# Optical metabolic imaging of heterogeneous drug response in pancreatic cancer patient organoids

**DOI:** 10.1101/542167

**Authors:** Joe T. Sharick, Christine M. Walsh, Carley M. Sprackling, Cheri A. Pasch, Alexander A. Parikh, Kristina A. Matkowskyj, Dustin A. Deming, Melissa C. Skala

**Affiliations:** Department of Biomedical Engineering, Vanderbilt University, Nashville, TN; Morgridge Institute for Research, Madison, WI; University of Wisconsin Carbone Cancer Center, Madison, WI; Division of Oncologic and Thoracic Surgery, University of South Carolina School of Medicine – Greenville, Greenville, SC; Department of Pathology and Laboratory Medicine, University of Wisconsin, Madison, WI; William S. Middleton Memorial Veterans Hospital, Madison, WI; Division of Hematology and Oncology, Department of Medicine, University of Wisconsin, Madison, WI; McArdle Laboratory for Cancer Research, Department of Oncology, University of Wisconsin, Madison, WI; Department of Biomedical Engineering, University of Wisconsin, Madison, WI

## Abstract

New tools are needed to match pancreatic cancer patients with effective treatments. Patient-derived organoids offer a high-throughput platform to personalize treatments and discover novel therapies. Currently, methods to evaluate drug response in organoids are limited because they cannot be completed in a clinically relevant time frame, only evaluate response at one time point, and most importantly, overlook cellular heterogeneity. In this study, non-invasive optical metabolic imaging (OMI) of cellular heterogeneity in organoids was evaluated as a predictor of clinical treatment response. Organoids were generated from fresh patient tissue samples acquired during surgery and treated with the same drugs as the patient’s prescribed adjuvant treatment. OMI measurements of heterogeneity in response to this treatment were compared to later patient response, specifically to the time to recurrence following surgery. OMI was sensitive to patient-specific treatment response in as little as 24 hours. OMI distinguished subpopulations of cells with divergent and dynamic responses to treatment in living organoids without the use of labels or dyes. OMI of organoids agreed with long-term therapeutic response in patients. With these capabilities, OMI could serve as a sensitive high-throughput tool to identify optimal therapies for individual pancreatic cancer patients, and to develop new effective therapies that address cellular heterogeneity in pancreatic cancer.

**Significance:** OMI can non-invasively quantify cellular-level heterogeneity in treatment response within pancreatic cancer patient-derived organoids, which could enable high-throughput drug screens to personalize treatment for individual patients and to accelerate drug discovery.

## Introduction

Pancreatic cancer has one of the lowest 5-year survival rates of all cancer types (8%) (1), and is projected to be the second leading cause of cancer death by 2030 (2). Surgery is required to cure pancreatic cancer (3), but can only be performed in 5-10% of cases, and only offers a 10.4% 5-year survival rate (4). Chemotherapy can enhance survival after surgery and standard treatments in the adjuvant setting include gemcitabine and gemcitabine combined with capecitabine (5-fluorouracil (5-FU) pro-drug) (5). Promising treatments for early-stage pancreatic cancer following surgery include gemcitabine combined with nab-paclitaxel, and FOLFIRINOX. Currently, oncologists must weigh these drug treatment options for individual patients based solely on potential side effects and have no *a priori* indication of whether a pancreatic cancer patient will respond to standard therapies. There is a need for a predictive tool to screen and identify alternative therapies that would benefit individual patients with either resectable or more advanced unresectable pancreatic cancer. Such a tool would minimize the needless costs and side effects accompanying ineffective treatment, while maximizing long-term survival rates. Additionally, the discovery of novel therapies would be accelerated by the ability to rapidly investigate the clinical efficacy of many drug candidates for many patients.

Tailoring treatment based on genomic analysis alone is insufficient due to poor understanding of the connections between pancreatic tumor driver mutations and drug response. Despite being less genetically diverse than most cancers, targeted approaches to treating pancreatic cancer have failed in clinical trials and drug treatment is limited to cytotoxic chemotherapies. Most pancreatic tumors do not express a specific therapeutic target (6), and the driver mutations present in a patient’s cancer do not predict their response to chemotherapy. Alternatively, pancreatic organoids have emerged as an appealing method to tailor treatments by performing high-throughput drug screening directly on a patient’s tumor cells (7-9). *In vitro* organoids recapitulate the genetic and histopathological characteristics of the original pancreatic tumor, along with its complex 3-dimensional organization (10-14). Organoid cultures also preserve interactions between tumor cells, immune cells (15), and fibroblasts (16), which can influence tumor drug response and are potential drug targets (17, 18). Generally, methods for measuring drug response in organoids have involved either cell viability assays, pooling of proteins, DNA, and RNA from many organoids, or tracking of organoid diameter changes. These methods homogenize the response of an entire organoid or many organoids and ignore cellular heterogeneity, which drives tumor treatment resistance (19-22). It is possible that minority subpopulations of lethal drug-resistant cells go completely undetected without more advanced assessment tools. Additionally, these methods generally neglect cellular metabolism, which is a major factor determining cellular drug response and heterogeneity (23-25).

Optical metabolic imaging (OMI) is a non-destructive, high-resolution fluorescence microscopy technique that quantifies the metabolic state of individual cells within a single organoid using cellular autofluorescence (26, 27). The fluorescence properties of NADH and NADPH overlap and are referred to as NAD(P)H. NAD(P)H, an electron donor, and FAD, an electron acceptor, are fluorescent metabolic co-enzymes present in all living cells. The optical redox ratio, defined as the ratio of the fluorescence intensity of NAD(P)H to that of FAD, reflects the redox state of the cell (28-30), and is sensitive to shifts in metabolic pathways (27, 31, 32). The fluorescence lifetimes of NAD(P)H and FAD are both two-exponential with distinct lifetimes for the free-and protein-bound conformations, and thus reflect the protein-binding activities of NAD(P)H and FAD (33-35). The optical redox ratio, NAD(P)H, and FAD fluorescence lifetimes all provide complimentary information, and can be combined into a composite endpoint called the OMI index (36). This metric distinguishes drug-resistant and responsive cells by their metabolic states and is robust and sensitive in pancreatic organoids (7).

OMI of organoids could improve predictions of patient outcomes for several reasons. First, drug-induced changes in cell metabolism precede changes in tumor size or overall cell viability (7, 27, 36, 37), and thus can measure drug response faster than conventional methods such as apoptosis and proliferation assays. Second, OMI analysis of cell subpopulations identifies and quantifies tumor heterogeneity (37, 38), which is vital to accurately capture patient drug response. Finally, OMI is non-invasive and does not require exogenous labels, so treatment response can be tracked over time in the same organoids. This is not possible with standard techniques which, by necessity, destroy samples. Therefore, OMI provides a fast, dynamic method to evaluate heterogeneous drug response at the organoid and single-cell level, integrating tumor heterogeneity into treatment planning and drug discovery. OMI of organoids has been validated as an accurate predictor of *in vivo* drug response in mouse models of pancreatic cancer (7), but has not yet been evaluated in humans. This study is the first to demonstrate that OMI of cellular metabolic heterogeneity in pancreatic tumor organoids provide an early measure of long-term *in vivo* drug response for individual patients.

## Materials and methods

### Tissue processing and organoid culture

Human tissue was collected with informed consent from all patients, and all studies were approved by the Institutional Review Board at the University of Wisconsin-Madison. Surgically resected tissue was placed in cold chelation buffer on ice for one hour. Tissues were washed with PBS and digested at 37°C in DMEM/F12 medium (Invitrogen) containing 1 mg/mL collagenase (Sigma), 0.125 mg/mL dispase (Invitrogen), 10% FBS (Gibco), and 1% pen-strep (Gibco) for 2-3 hours with intermittent shaking. The resulting cell macro-suspension was rinsed in PBS, re-suspended in 1:1 DMEM/F12:Matrigel, plated in 50 μl droplets, and allowed to solidify at 37°C, 5% CO_2_ in a cell incubator. Once solidified, droplets were overlaid with DMEM/F12 supplemented with 7% FBS, 20 μM Y-27632 (Sigma), 50 ng/ml EGF (Invitrogen), RSPO-conditioned medium (homemade) and 1% penicillin-streptomycin (Gibco). FBS, Y-27632, and RSPO-conditioned medium were removed from cultures if fibroblasts were out-growing tumor cells.

### Drug screening

24 hours prior to imaging, media was replaced with fresh media containing 85 μM gemcitabine (39), 10 μM 5-FU, 200 nM TAK-228 (40, 41), 250 nM ABT-263 (42, 43), 5 μM oxaliplatin, 10 μM nab-paclitaxel, 50 nM SN-38 or combinations of each. Doses were selected to replicate clinically relevant peak plasma concentrations. FOLFIRINOX treatment was comprised of 5-FU, oxaliplatin, and SN-38. After the first imaging time point (24 hours), gemcitabine, nab-paclitaxel, SN-38, and oxaliplatin were removed from cultures to simulate single bolus delivery, while 5-FU, TAK-228, and ABT-263 exposure was maintained throughout the experiment (to simulate daily oral delivery). Chemotherapy drugs were obtained from the University of Wisconsin Carbone Cancer Center Pharmacy. TAK-228 was obtained from LC Laboratories, and ABT-263 was obtained from Apex Bio.

### Multiphoton imaging

Fluorescence imaging was performed using a custom multiphoton fluorescence lifetime system (Bruker Fluorescence Microscopy). A 40x water immersion objective (Nikon, 1.15 NA) was used with an inverted microscope (Nikon, TiE). A titanium:sapphire laser (Spectra-Physics InSight DS+) was used for excitation, while GaAsP photomultiplier tubes (H7422P-40, Hamamatsu) detected emission light. 750 nm and 890 nm light were used for two-photon excitation of NAD(P)H and FAD, respectively. A 440/80 nm filter was used to collect NAD(P)H fluorescence emission, and a 550/100 nm filter was used to collect FAD fluorescence emission. Images were acquired over 60 seconds, with a pixel dwell time of 4.8 μs for 256×256 pixel images. Fluorescence lifetime data with 256 time bins was acquired using time-correlated single photon counting electronics (SPC-150, Becker & Hickl). A Fluoresbrite YG microsphere (Polysciences) was imaged daily as a fluorescence lifetime standard, which had a stable lifetime (2.09 ± 0.05 ns, n=61), consistent with previously published values (26, 33, 35).

### Organoid imaging

Imaging of organoids was performed in 35 mm glass-bottom dishes (#P35G-1.5-14-C, MatTek). At least five representative organoids were imaged in each treatment group at each time point. At least two additional images of the fibroblast monolayer on the coverslip were taken, if present. Images were acquired 1, 2, 3, 5, and 7 days after initial treatment.

### Cyanide experiment

Four previously untreated organoids from Patient 1 underwent OMI immediately before and after the addition of media containing 12 mM NaCN (Sigma), for a final concentration of 6 mM. OMI endpoints were quantified at the single-cell level, at least 60 cells per group. Redox ratio values were normalized to the pre-treatment average.

### Image analysis

NAD(P)H and FAD fluorescence lifetime images were analyzed using SPCImage software (Becker & Hickl) (44). Briefly, a histogram of photon counts per temporal bin, or decay curve, is generated for each pixel by binning the photon counts of all 9 surrounding pixels. This decay curve is deconvolved with the instrument response function, and then fit to a two-component exponential decay (Equation 1).

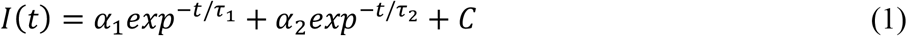

Here, *I*(*t*) represents the fluorescence intensity measured at time *t, α*_1_ and *α*_2_ represent the fractional contributions of the short and long lifetime components to the overall signal respectively, *τ*_1_ and *τ*_2_ are the short and long lifetime components respectively, and *C* represents background light. The two lifetime components are used to distinguish between the free and bound forms of NAD(P)H and FAD (45, 46). The mean lifetime (*τ*_*m*_) is a weighted average of the free and bound lifetimes, and is calculated for both NAD(P)H and FAD in each pixel using Equation 2.

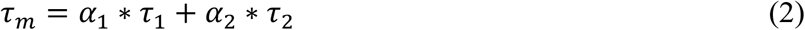

The optical redox ratio was calculated for each pixel by dividing the intensity of NAD(P)H by the intensity of FAD. A customized CellProfiler (v1.0.6025) routine was written to mask individual cell cytoplasms and extract average NAD(P)H and FAD intensities and lifetime components for each cell cytoplasm (47, 48). All reported redox ratios are normalized to average control values of the same patient and time point.

### OMI index calculation

The OMI index, in this study, is a linear combination of three independent OMI endpoints (redox ratio, NAD(P)H *τ*_*m*_, and FAD *τ*_*m*_), each centered around the average value measured in control cells within each patient at the same experimental time point. This differs from previous descriptions (7, 36) where the end points are mean-centered across all cells in all treatment groups. This modification allows drug responses to be compared between patients in this study. As before, the redox ratio, NAD(P)H *τ*_*m*_, and FAD *τ*_*m*_ are given coefficients of (1,1,-1). A decrease in OMI index relative to control correlates with drug response, while an increase or lack of change indicates drug resistance.

### Subpopulation analysis

A Gaussian mixture distribution model was used to assess heterogeneity of cellular metabolism (7, 36, 38, 49). OMI values for all cells within a treatment group and time point are input into this model described by Equation 3.

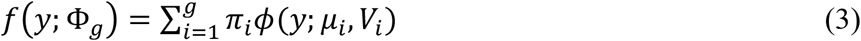

Here, *g* represents the number of subpopulations in the model, *ϕ*(*y*; *μ*_*i*_, *V*_*i*_) is a normal probability density function where *μ*_*i*_ represents the mean and *V*_*i*_ represents variance, and *π*_*i*_ is the mixing proportion. Models containing *g* = 1, 2, and 3 subpopulations are fit to the data, and the goodness of fit for each model is assessed using the Akaike information criterion (AIC) (50). The best fit of the three models, equivalent to the lowest AIC, is used to evaluate heterogeneity. For comparison, distributions are normalized such that all have an area under the curve of 1. The weighted heterogeneity index (wH-index, Eq. 4) is based on the Gaussian distribution models described by Eq. 3, and is a modified form of the Shannon diversity index used to quantify the degree of heterogeneity (37, 51).

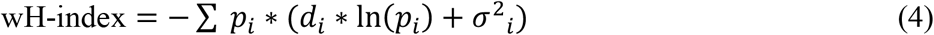

Here, *i* represents each subpopulation in the Gaussian distribution model, *d* represents the distance between the median of each subpopulation and the median of the entire distribution, *p* represents the proportion of all cells belonging to that subpopulation, and *σ*^2^ is the variance.

### Organoid immunofluorescence

Organoids were rinsed in PBS and fixed for 20 minutes in 4% paraformaldehyde (VWR). Fixed organoids were rinsed and stored in PBS at 4°C until stained. Organoids were blocked for one hour in PBS with 10% goat serum and 0.3% Triton-X 100 (Sigma) at room temperature followed by co-incubation with Ki67 antibody conjugated to AlexaFluor 488 (1:50, Cell Signaling #11882S) and CC3 antibody conjugated to AlexaFluor 555 (1:50, Cell Signaling #9604S). Organoids were then washed in PBS and mounted to a slide using ProLong™ Diamond Antifade Mountant with DAPI (Molecular Probes). DAPI was imaged on the multiphoton microscope at 40x magnification using 750 nm for excitation and 440/80 nm bandpass filter for emission. AlexaFluor 488 was imaged using 965 nm for excitation and 550/100 nm filter for emission. AlexaFluor 555 was imaged using 1050 nm excitation and a 585/65 nm filter. Six or more organoids per treatment group were imaged and the percentage of Ki67-positive cells and cleaved caspase-3 positive cells in each organoid were quantified.

### Patient-derived xenografts

Animal research was approved by the UW-Madison Institutional Animal Care and Use Committee. Organoids from Patient 13 were pelleted, re-suspended in media, and mixed 1:1 with Matrigel. This mixture was subsequently injected (100 μl) subcutaneously into bilateral flanks of female NOD *scid* gamma mice at 6 weeks old (NOD.Cg-*Prkdc*^*scid*^ *Il2rg*^*tm1Wj1*^/SzJ, The Jackson Laboratory) for initial PDX establishment. For treatment experiments, extracted tumors were mechanically minced to form a cell suspension, which was then mixed with Matrigel for injection into experimentally naïve female athymic nude mice at 6 weeks old (Hsd:Athymic nude-Foxn1^nu/nu^, Envigo). Tumor volume was measured with calipers using the formula 0.5*length*width^2^. When average tumor volume reached ∼150 mm^3^, mice were randomized into two groups. 20 mice received 100 mg/kg gemcitabine and 100 mg/kg nab-paclitaxel weekly via intraperitoneal injection while 23 control mice received only PBS weekly. Tumor volume was measured twice weekly. Mice were euthanized and tumors were collected when humane endpoints were reached.

### High-depth targeted gene sequencing

Patient 13 organoids were sequenced using the Qiagen Comprehensive Cancer Panel and molecular barcode technology, with greater than 500x median coverage.

### UMAP clustering

UMAP (Uniform Manifold Approximation and Projection) was performed on metabolic imaging data from all organoid cells (control and treated) from all patients and time points (N ∼ 45,000) using the UMAP.py package (52). Nine metabolic measures: redox ratio, NAD(P)H *τ*_*m*_, NAD(P)H *α*_1_, NAD(P)H *τ*_1_, NAD(P)H *τ*_2_, FAD *τ*_*m*_, FAD *τ*_1_, FAD *τ*_2_, and FAD *α*_1_ were reduced to two components. Points could have 10 neighbors in the manifold structure, and cosine was used to measure distance in the multivariate space. The two resulting components were visually displayed using the CRAN package ggplots2 (53).

### Statistics

Differences in OMI index, optical redox ratio, NAD(P)H *τ*_*m*_, FAD *τ*_*m*_, CC3+%, and Ki67+% between treatment groups were tested using a Wilcoxon rank-sum test. Normalized tumor volumes were compared using a student t-test and a D’agostino-Pearson normality test. Treatment effect size was calculated with Glass’s Δ (54).

## Results

### Organoid generation, drug screening, and optical metabolic imaging

Organoids were generated from fresh patient tissue samples acquired during distal pancreatectomy or pancreaticoduodenectomy (Whipple resection) surgeries. The overall rate of successful organoid formation was 67% (12 of 18 patients) (Fig. 1A), including mostly pancreatic tumors (pancreatic ductal adenocarcinomas (PDAC) and anaplastic carcinoma of the pancreas), along with two pancreatic intraepithelial neoplasia (PanIN) lesions and one ampullary adenocarcinoma (Supplementary Table S1). 50% of the successfully cultured patient samples underwent neoadjuvant treatment prior to resection, including one patient that was downgraded from PDAC to PanIN following a complete pathologic response to chemotherapy (Patient 5). Neoadjuvant treatment did not impede organoid formation (75% and 60% success rates for pretreated and non-pretreated samples respectively). Viability and OMI were validated for pancreatic organoids by quantifying metabolic inhibition to cyanide, a known inhibitor of the electron transport chain (Supplementary Fig. S1). The effects of cyanide on OMI endpoints agreed with previous reports (27). After an establishment period, organoids were treated with a panel of standard and experimental pancreatic cancer therapies and imaged over a time course (Fig. 1B). This drug panel included 5-FU and gemcitabine chemotherapy, along with an experimental combination of TAK-228 (mTORC1/2 inhibitor) and ABT-263 (Bcl-2 and Bcl-xL inhibitor). Strategies to inhibit mTOR kinase activity could potentially induce apoptosis (55, 56), but this process is inhibited by antiapoptotic proteins of the Bcl-2 family (57). Thus, the combination of an mTORC1/2 inhibitor (TAK-228) with a Bcl-2 and Bcl-xL inhibitor (ABT-263) could strongly induce apoptosis in pancreatic cancer. Additional standard drugs were tested on organoids from Patients 6, 13, and 18 at later time points after patient treatment plans were obtained. The length of the organoid establishment period varied by patient between 4 and 12 days (Fig. 1C), and ended when organoids were clearly visible and proliferating with rounded edges. Differences in size and cellular quality of patient tumor samples likely contributed to the variance in time to maturation for organoid lines. One line (Patient 6) was thawed after being frozen, requiring additional time to re-establish a usable culture. Representative images demonstrate that multiphoton microscopy measures OMI endpoints with high resolution (Fig. 1D-F). This allows endpoints to be quantified in individual cells by masking each cell nucleus and cytoplasm using NAD(P)H fluorescence intensity (Fig. 1G). Tumor cells often grew as 3-dimensional hollow spheres (Fig. 1H), but other morphologies such as solid spheres were noted. The effects of drugs on the three OMI endpoints at 72 hours is summarized by comparing the mean values across all cells (Fig. 1I-K). A decrease in the mean optical redox ratio or NAD(P)H *τ*_*m*_ (see Equation 2), or an increase in the mean FAD *τ*_*m*_ is considered an indicator of response to that treatment, based on previous studies in breast organoids (36). The OMI index combines the complimentary information provided by the three endpoints for each individual cell (Fig. 1L), and a decrease in the mean OMI index with treatment indicates a metabolic response. At the cellular level, Patient 14 organoids did not respond to 5-FU on average, but did respond to gemcitabine and targeted therapy (all p<0.05 vs. control).

**Figure 1.**
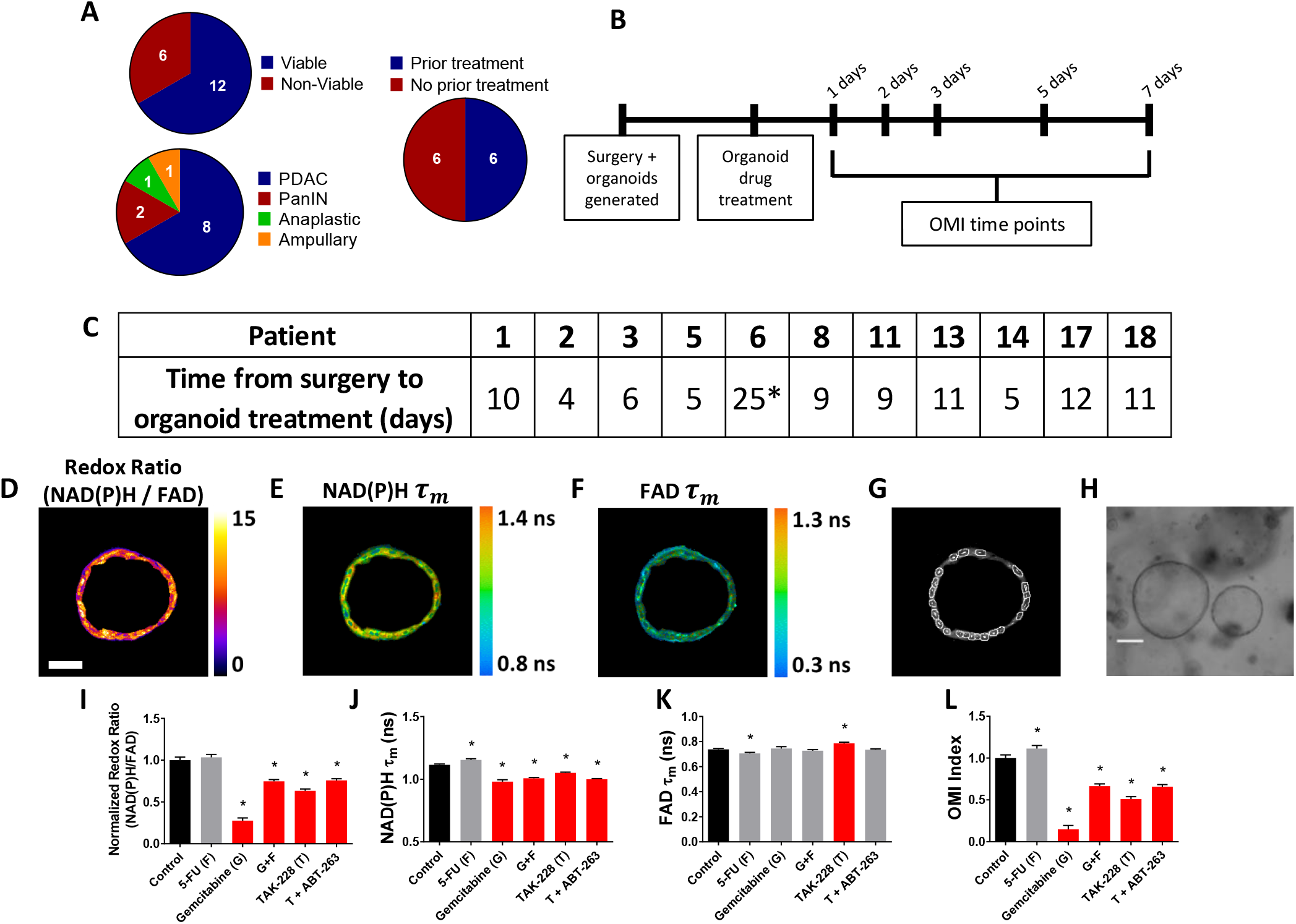
Pancreatic organoid generation, drug screening, and optical metabolic imaging. **A,** Pie charts depicting the success rate for generating viable organoids from patient pancreatic lesions (top left), the distribution of PDAC, PanIN, anaplastic cancer, and ampullary cancer among successfully generated organoids (bottom left), and the distribution of previously treated versus untreated tumors among successfully generated organoids (right). **B,** Experimental workflow begins with a brief period where organoids are allowed to expand after initial generation, followed by drug treatment, and a series of time points where OMI is performed to track dynamic organoid drug responses. **C,** Time between organoid generation and drug treatment for each individual patient in the study. * Patient 6 organoids were thawed from frozen stocks and required additional time in culture. Representative redox ratio (**D**), NAD(P)H *τ*_*m*_ (**E**), and FAD *τ*_*m*_ (**F**) images of an untreated pancreatic organoid taken 6 days after surgical resection (Patient 14). Scale bar is 50 μm. **G**, Masks of individual cell cytoplasms overlaid onto NAD(P)H intensity image. **H,** Representative brightfield image of pancreatic organoids (Patient 14). Scale bar is 200 μm. The effect of 72 hours of drug treatment on the redox ratio (**I**), NAD(P)H *τ*_*m*_ (**J**), and FAD *τ*_*m*_ (**K**) of pancreatic organoids quantified at the single-cell level (Patient 14). **L,** The effect of 72 hours of drug treatment on OMI index, a composite metric of metabolic drug response (Patient 14). Error bars indicate mean ± SEM. * p<0.05 vs. control. Red bar indicates response to treatment (significant reduction in mean redox ratio, NAD(P)H *τ*_*m*_, or OMI index, or significant increase in mean FAD *τ*_*m*_).

### OMI of organoids resolves differential sensitivities to relevant drug treatments

A wide variety of OMI index responses were elicited across treatment conditions and patient samples. In addition to statistical significance of these drug responses, treatment effect sizes were calculated on all OMI measurements in order to determine their magnitude. A heatmap of OMI index treatment effect size, calculated using Glass’s Δ at 72 hours post-treatment, shows significant inter-patient heterogeneity for drug response in organoids (Fig. 2A; additional drugs and time points in Supplementary Fig. S2). In a subset of patient samples, a significant fibroblast population (co-cultured with organoids) migrated from the 3D matrix and adhered to the 2D glass coverslip. Heterogeneity in these fibroblasts has been shown in organoid models of murine pancreatic cancer (7, 58). A heatmap of OMI index treatment effect size, calculated using Glass’s Δ at 72 hours post-treatment, also shows inter-patient drug response heterogeneity in co-cultured fibroblasts (Fig. 2B; Supplementary Fig. S3). Not all drugs were tested on every patient’s cells for the full time-course. Drug choices were made based on the availability of viable organoids and the clinical treatment plans (or lack thereof) for individual patients. All organoids were initially treated with the drug panel in Figure 1 because gemcitabine and 5-FU combination therapy was most likely to be prescribed after surgery (excluding Patient 6; it was known in advance that they would receive oxaliplatin rather than gemcitabine). When additional drugs were prescribed for a patient such as nab-paclitaxel or the FOLFIRINOX regimen, their panels were expanded to incorporate those treatments. Responses to a chemotherapy drug (5-FU) and an experimental targeted therapy combination (TAK-228 and ABT-263) at days 1, 3, and 7 demonstrate how OMI can track single-cell drug responses over time within organoids (Fig. 2C; additional drugs and time points in Supplementary Fig. S4) and co-cultured fibroblasts (Fig. 2D; Supplementary Fig. S5). For example, response data show an increased response to TAK-228 and ABT-263 combination targeted therapy over time, while 5-FU remained ineffective for most patients.

**Figure 2.**
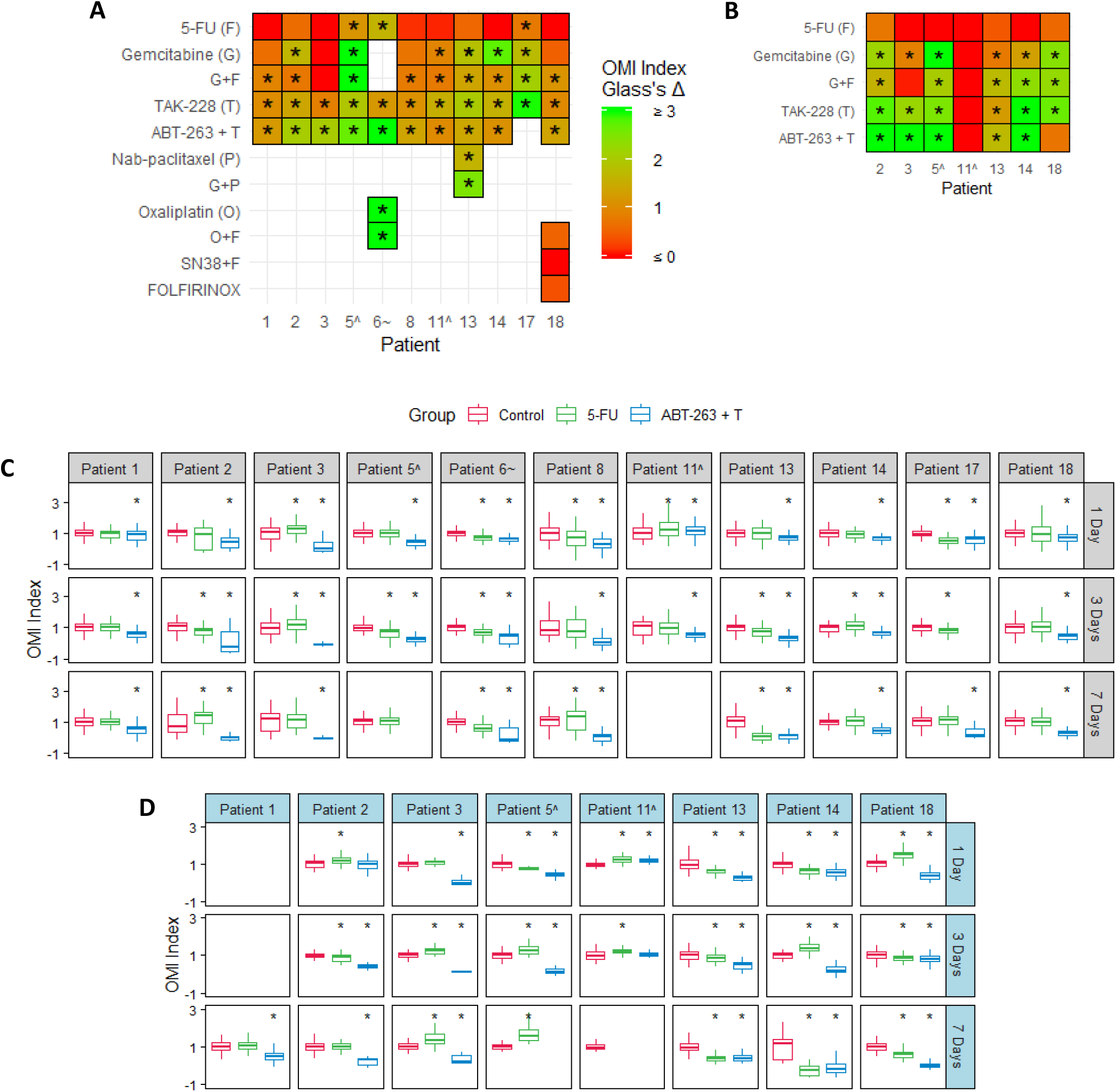
OMI of organoids resolves differential sensitivities to relevant drug treatments. **A** and **B,** Heatmap representation of the OMI index treatment effect size (Glass’s Δ) at 72 hours in organoids (**A**) and fibroblasts co-cultured with organoids (**B**). * Glass’s Δ ≥ 0.75. **C** and **D,** Boxplot comparing metabolic responses to 5-FU and ABT-263 combined with TAK-228 (A+T) between patients and time points in organoids (**C**) and fibroblasts co-cultured with organoids (**D**). Response is measured using OMI index at the single-cell level. A decrease in OMI index versus control indicates response to treatment. Center line indicates median, box indicates interquartile range (IQR), and whiskers indicate most extreme data point within 1.5*IQR. ‘^’ and ‘∼’ indicate the patient lesion was diagnosed as PanIN or ampullary cancer, respectively. * p<0.05 vs. control.

### OMI captures non-genetic cellular heterogeneity in pancreatic organoids

Additional analysis was performed on Patient 13 organoids to evaluate the utility of OMI. The combination of gemcitabine and nab-paclitaxel was used in addition to the standard treatment panel on Patient 13’s organoids to mimic the treatment received prior to sample collection (Fig. 3A). Population density modeling was used to determine whether cellular subpopulations of metabolic response were present in organoids for each treatment condition at 72 hours (Fig. 3B; Supplementary Fig. S6A). Metabolic subpopulations were observed in controls and organoids treated with gemcitabine and nab-paclitaxel (G+P) combination. Patient 13 responded poorly to this treatment prior to surgery and organoid generation. OMI of organoids treated with the experimental combination of TAK-228 and ABT-263 resulted in a single, homogeneous drug-responsive population. Both treatments reduced the degree of heterogeneity present in the organoids, measured with the wH-index, but this reduction was larger with TAK-228 + ABT-263 treatment compared to the standard chemotherapies that the patient received (Fig. 3C; additional drugs in Supplementary Fig. S6B).

**Figure 3.**
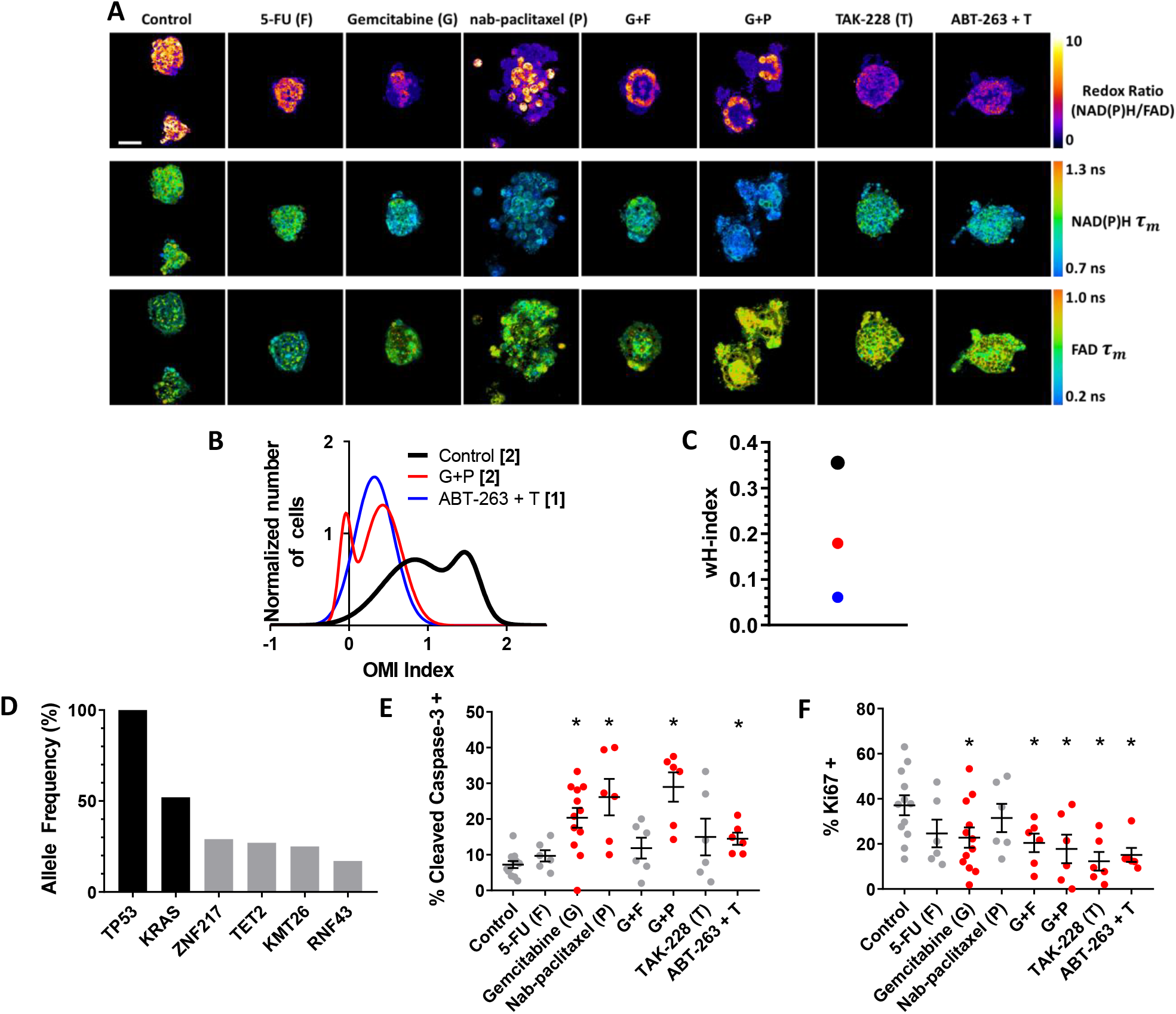
OMI captures non-genetic cellular heterogeneity in pancreatic organoids. **A,** Representative images of the redox ratio, NAD(P)H *τ*_*m*_, and FAD *τ*_*m*_ in organoids generated from Patient 13 (anaplastic carcinoma of the pancreas), treated with standard chemotherapies and experimental targeted therapies for 72 hours. Scale bar is 50 μm. **B,** Normalized density distributions of the OMI index of individual cells contain subpopulations with G+P treatment, but not ABT-263 +T treatment after 72 hours. Bracketed number indicates number of subpopulations. **C,** The effect of treatment on the wH-index of OMI index density distributions in (B). Error bars not visible. N=1000 fits/group. **D,** High-depth targeted cancer gene sequencing of Patient 13 organoids. Allele frequencies of ∼50% for *KRAS* and 100% for *TP53* were found (black bars). Alterations with allele frequencies of 10-30% were detected (gray bars), but none of these alterations were pathogenic. **E,** Cleaved caspase-3 staining of organoids shows differences in apoptosis between treatment conditions after 72 hours of treatment. **F,** Ki67 staining of organoids shows differences in proliferation between treatment conditions after 72 hours. Each dot represents one organoid (mean ± SEM), and red indicates significant response to treatment. *p<0.05 vs. control.

High-depth targeted gene sequencing was performed on untreated Patient 13 organoids to determine whether subclonal populations could be resolved to explain the metabolic heterogeneity (Fig. 3D). A mutation in *TP53* tumor-suppressor gene (stop-gain Gln165*) and a mutation in the *KRAS* oncogene (G12V) were found with allele frequencies of 100% and 52%, respectively, indicating a single population of cells with homogeneous driver mutations. Mutations with allele frequencies between 10-30% are indicative of potential subclonal populations (59). Only 4 alterations were found to occur within this range, with 3 of the 4 at frequencies just below the top of this range. None of the alterations identified are pathogenic or known to alter tumor biology. This indicates that this patient sample was quite homogeneous on a genetic level and the treatment subpopulations identified are likely related to metabolic changes and not differing mutation profiles. To test whether OMI agreed with traditional cell markers, a subset of organoids were fixed at 72 hours of treatment and dual-stained using immunofluorescence for cleaved caspase-3 (CC3) and Ki67 to quantify apoptosis and proliferation rates, respectively (Fig. 3E,F; Supplementary Fig. S7). Three treatments caused both a significant increase (p<0.05 vs. control) in apoptosis and a significant decrease in proliferation: gemcitabine, gemcitabine and nab-paclitaxel, and TAK-228 and ABT-263. When averages were compared, proliferation rates correlated with changes in cell metabolism measured by OMI index (p<0.001), but apoptosis rates did not (Supplementary Fig. S8). A patient-derived xenograft line was generated and athymic nude mice were treated with combination therapy of gemcitabine and nab-paclitaxel (Supplementary Fig. S9) to determine if *in vivo* treatment response correlates with that seen in organoids. Tumor growth was tracked by direct caliper measurement and an early reduction was observed over the first 7 days (p<0.05); however, the effect was not sustained.

### Heterogeneity of drug response in organoids agrees with later patient recurrence during adjuvant therapy

Seven patients received adjuvant drug treatment following surgical resection of their tumor. Clinical treatment efficacy was tracked and compared to the OMI prediction of drug response at 72 hours post-treatment in patient-matched organoids. Four patients whose organoids exhibited a homogeneous response to the patient’s prescribed therapy were classified as “predicted responders” (Fig. 4). Representative redox ratio, NAD(P)H *τ*_*m*_, and FAD *τ*_*m*_ images demonstrate the morphological variety as well as a decrease in size or structural integrity with treatment of organoids generated from these patients (1, 2, 6, and 14) (Fig. 4A-D). Representative images of collapsed organoids that lose their hollow morphology (Fig. 4C,D), are the result of treatment sensitivity and are generally not present in untreated cultures. Furthermore, the OMI index reflects an organoid response to the prescribed treatment for these patients (p<0.0001) (Fig. 4E-H). Glass’s Δ effect size was calculated for each treatment’s OMI index value in addition to statistical significance, and was found to be consistently greater than 0.75 for these patients. Cellular population density modeling of organoids from Patients 1, 2, 6, and 14 did not reveal distinct metabolic subpopulations of response (Fig. 4I-L). In each case, one homogeneous population of response was found. Fibroblasts co-cultured with organoids from Patients 2 and 14 also showed homogeneous response to the patient’s prescribed therapy (Supplementary Fig. S10).

**Figure 4.**
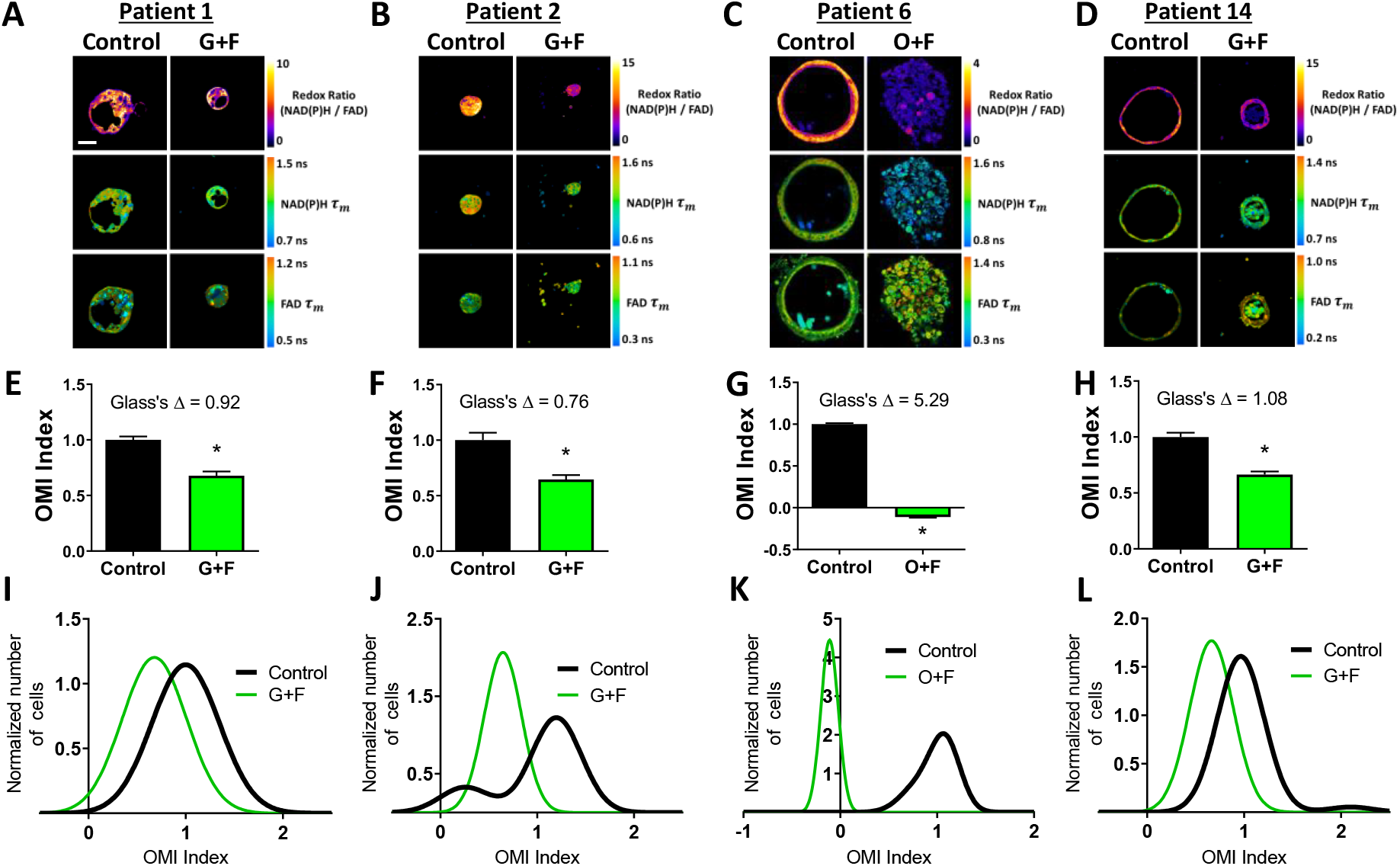
Organoids of Patients 1, 2, 6, and 14 exhibit homogeneous response to patient therapy. **A-D,** Representative redox ratio, NAD(P)H *τ*_*m*_, and FAD *τ*_*m*_ images of pancreatic organoids from Patients 1 (**A**), 2 (**B**), 6 (**C**), and 14 (**D**). Left columns indicate control organoids, and right columns indicate organoids treated with the same drugs as the patient adjuvant treatment. Scale bar is 50 μm. **E-H,** The effect of the same drugs on the OMI index averaged across all cells in organoids derived from Patient 1 (**E**), 2 (**F**), 6 (**G**), and 14 (**H**). Error bars indicate mean ± SEM. * p<0.0001. **I-L,** Single-cell OMI index subpopulation analysis of treatment response in organoids from Patient 1 (**I**), 2 (**J**), 6 (**K**), and 14 (**L**).

Three patients (3, 8, and 18) whose organoids exhibited treatment response heterogeneity were classified as “predicted non-responders” (Fig. 5). Representative images show a lack of apparent response in terms of OMI endpoints (Fig. 5A-C). On average, cells from Patient 3 and 8 organoids did not respond to the patient’s prescribed treatment (p<0.05 increase in OMI index and no significant change, respectively, Fig. 5D,E). Unlike 3 and 8, the OMI index of Patient 18’s organoid cells significantly decreased with treatment (p<0.0001, Fig. 5F), but further OMI analysis also predicts poor response for this patient. The treatment effect size was small (Glass’s Δ < 0.3) in organoids for all three patients, and all exhibited multiple subpopulations of tumor cells post-treatment, some of which overlapped completely with control distributions or contained OMI index values above control (Fig. 5G-I). Patient 3 organoids treated with gemcitabine and 5-FU exhibited a drug-responsive minority subpopulation with a smaller OMI index than the mean of the control population (yet still within the control distribution), but also a larger drug-resistant subpopulation that had a higher OMI index than the mean of the control population (Fig. 5G). Patient 8 organoids contained high and low OMI index subpopulations that were not affected by 5-FU treatment beyond a small increase in the percentage of cells falling in the high OMI index group (Fig. 5H). A majority of cells from Patient 18 treated with FOLFIRINOX showed response to treatment, but two FOLFIRINOX-resistant subpopulations remained (Fig. 5I). Fibroblasts co-cultured with organoids from Patient 3 also showed a lack of response to gemcitabine and 5-FU, along with treatment-induced metabolic heterogeneity (Supplementary Fig. S10).

**Figure 5.**
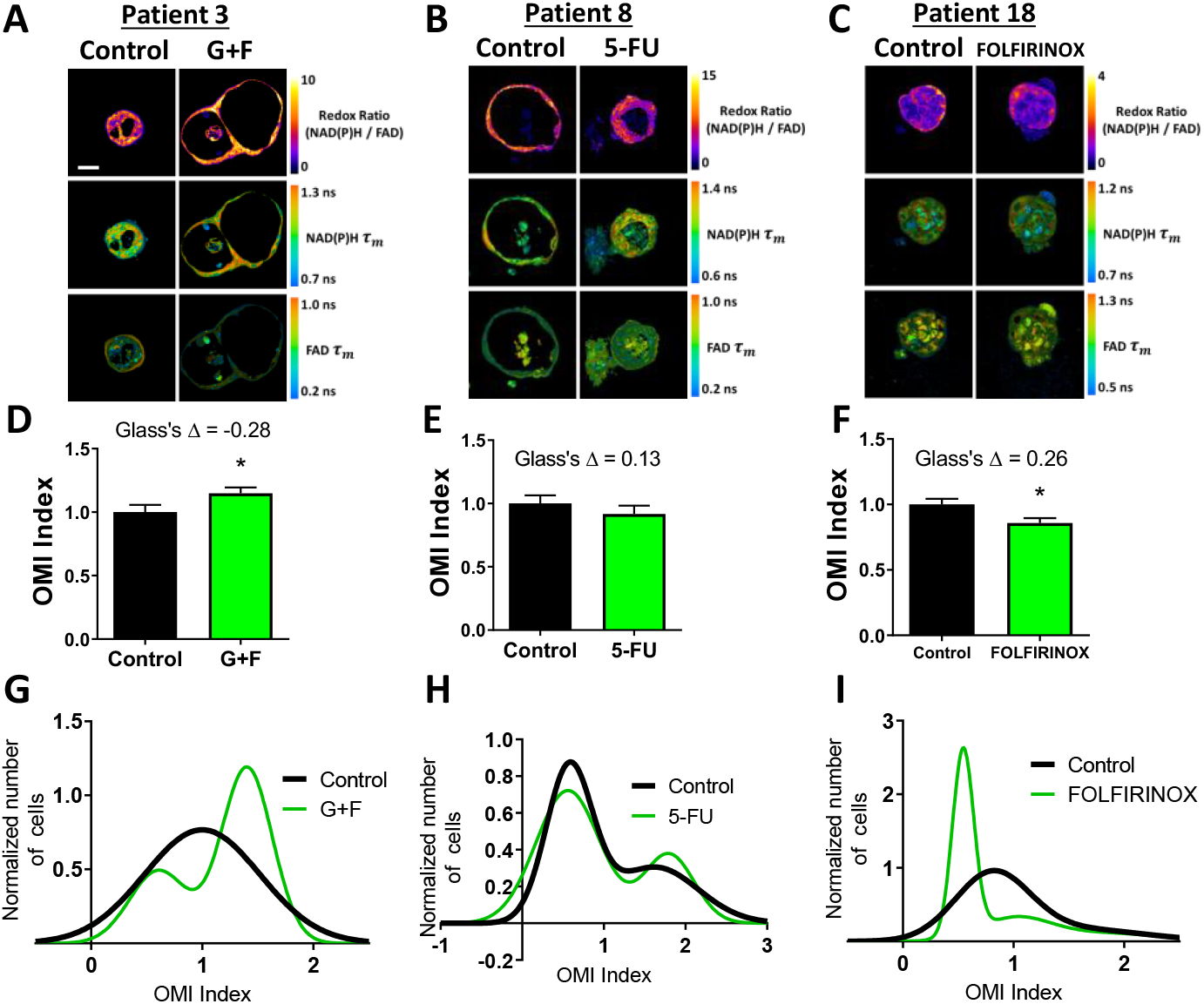
Organoids of Patients 3, 8, and 18 exhibit partial or no response to patient therapy. **A-C,** Representative redox ratio, NAD(P)H *τ*_*m*_, and FAD *τ*_*m*_ images of pancreatic organoids from Patients 3 (**A**), 8 (**B**), and 18 (**C**). Left columns indicate control organoids, and right columns indicate organoids treated with the same drugs as the patient adjuvant treatment. Scale bar is 50 μm. **D-F,** The effect of the same drugs on the OMI index averaged across all cells in organoids derived from Patient 3 (**D**), 8 (**E**), and 18 (**F**). Error bars indicate mean ± SEM. * p<0.05. **G-I,** Single-cell OMI index subpopulation analysis of treatment response in organoids from Patient 3 (**G**), 8 (**H**), and 18 (**I**).

The time between surgical resection and the first evidence of recurrence, or recurrence free survival (RFS) time, was plotted for these seven patients (Fig. 6A). One year of follow-up data is available for Patients 1, 2, 3, 6, 8, and 14. Patients 3 and 8, classified as predicted non-responders, experienced recurrences within one year. Patients 1, 2, 6, and 14, classified as predicted responders, each survived at least one year after surgery without a recurrence. One year of follow-up data is not yet available for 18. Patients 1, 6, 14, and 18 have no signs or symptoms of disease at the time of this writing. Patients with a RFS >12 months had a higher OMI index Glass’s Δ in organoids at 72 hours post-treatment than patients with a RFS <12 months (Fig. 6B). Single-cell OMI distributions in patients with a RFS >12 months were best fit with one subpopulation, while patients with a RFS <12 months were best fit with two (Fig. 6C). The degree of heterogeneity, quantified by the wH-index, decreased in treated vs. control organoids in patients with a RFS >12 months, and increased in treated vs. control organoids in patients with a RFS <12 months (Fig. 6D).

**Figure 6.**
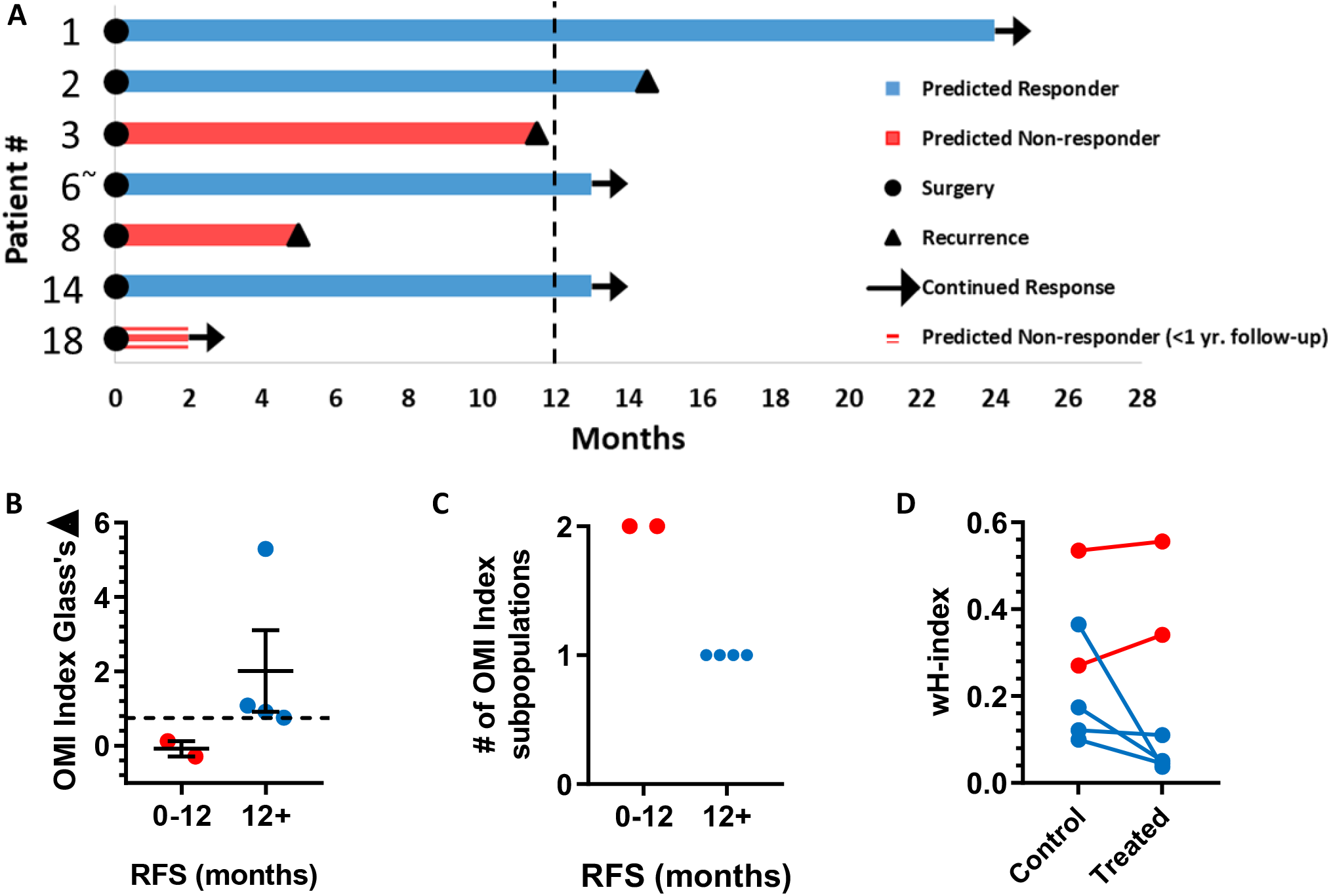
Patient clinical outcomes while on adjuvant therapy. **A,** Swimmer plot indicating the number of months without recurrence following surgical resection of the tumor and adjuvant treatment. Patients are classified as predicted responders and non-responders based on organoid response profiles. Arrows indicate that the patient continues to survive without recurrence at the time of publication. ‘^∼^’ indicates the patient’s lesion was diagnosed as ampullary cancer. **B,** Patients with RFS > 12 months had higher OMI index effect sizes at 72 hours (Glass’s Δ) than patients with RFS < 12 months (mean ± SEM). Dotted line represents proposed cutoff of Δ = 0.75. **C,** Patients with RFS > 12 months demonstrated one cell subpopulation in treated organoids by OMI index at 72 hours, while patients with RFS < 12 months showed two distinct subpopulations. **D,** Patients with RFS > 12 months show a decrease in wH-index with treatment compared to control organoids. Patients with RFS < 12 months show an increase in wH-index with treatment compare to control organoids. Error bars not visible. N=1000 fits/group.

### Dimensional reduction of single-cell OMI data reveals organoid cell clustering by patient of origin

UMAP dimensional reduction analysis was performed on the OMI variables of organoid cells imaged at all time points (45,752 cells) to determine whether cells from different patients exhibited unique metabolic profiles (Fig. 7). Cells from the same patient generally clustered together (Fig. 7A). Some patients (e.g. 6, 13, 17) formed a primary cluster and a secondary, smaller cluster. Plotting control and treated cells separately revealed that these secondary clusters occurred due to drug treatment (Fig. 7B,C).

**Figure 7.**
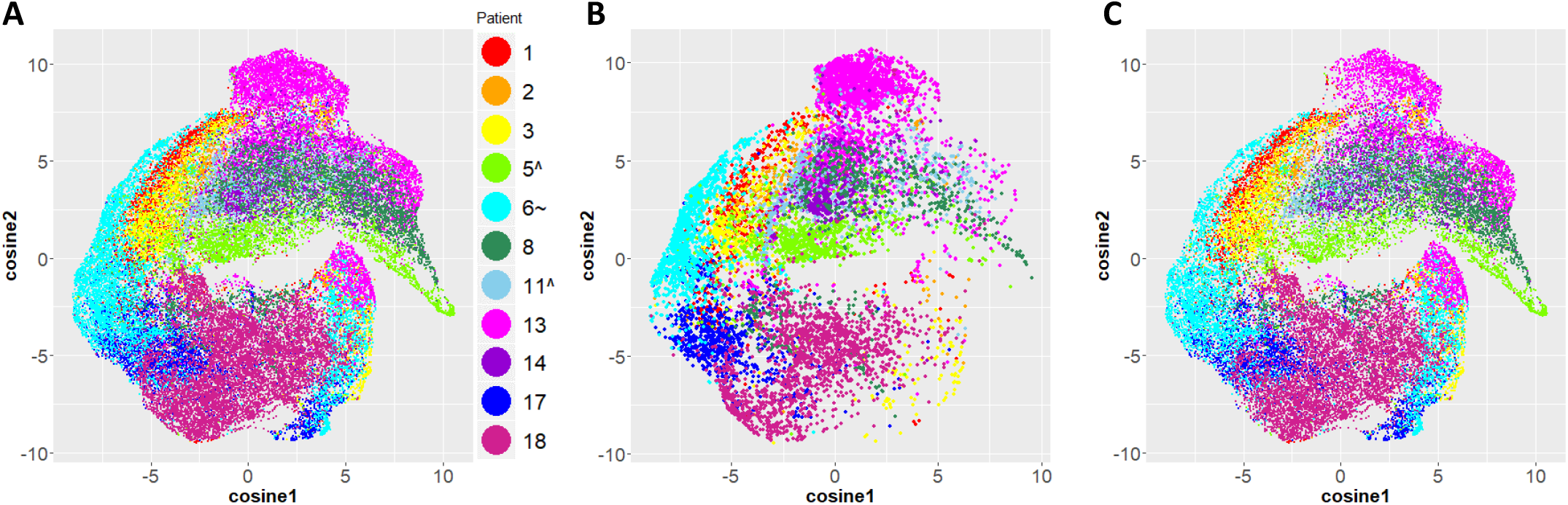
UMAP dimensional reduction analysis of organoid cell metabolism reveals clustering by patient. **A,** UMAP of all organoid cells from all time points color-coded post-analysis by patient of origin. **B,** Untreated cells only. **C,** Drug-treated cells only. ‘^’ and ‘∼’ indicate the patient lesion was diagnosed as PanIN or ampullary cancer, respectively.

## Discussion

Pancreatic organoids can be used for drug screens directly on patient cells, which could enable rational treatment planning for individual patients (7-9). Organoids also provide a platform to discover new drugs and drug combinations to treat pancreatic cancer patients, who currently suffer from a severe lack of effective treatment options. Existing methods to evaluate drug response in organoids ignore cellular heterogeneity, which can lead to patient relapse. Thus, our group developed OMI as a single-cell analysis tool to detect minority subpopulations of drug-resistant cells existing within living organoids that would otherwise appear responsive. We have previously shown that OMI detects subpopulations of drug response in murine PDAC organoids and early treatment responses in human pancreatic organoids (7). Here, we investigate for the first time whether OMI measurements of early drug response heterogeneity in organoids can capture meaningful treatment responses in individual pancreatic cancer patients.

Organoids were successfully generated from PDAC, precancerous PanIN, anaplastic carcinoma of the pancreas, and ampullary adenocarcinoma tissue samples, with equal success for both pre-treated and treatment-naïve samples. While some time is required to establish healthy organoids for drug screening, OMI measurements began within a 4-12 day window of the patient’s surgery. This is crucial for generating individualized treatment plans because pancreatic cancer is often advanced at diagnosis, and any delay to receiving effective therapy may increase mortality. Three independent OMI endpoints (redox ratio, NAD(P)H *τ*_*m*_, and FAD *τ*_*m*_) were quantified in response to panels of standard drugs (gemcitabine and 5-FU) and experimental targeted therapies that have shown promise in pancreatic cancer (TAK-228 and ABT-263) (Fig. 1). Each OMI endpoint captures unique metabolic information (27), and quantitatively combining these three independent measurements into the OMI index provides a technique to evaluate early metabolic response to treatment. An oncologist could use this technology to quickly determine if a patient would benefit from experimental targeted therapies over standard chemotherapies, rather than waiting for standard chemotherapy to fail while exposing the patient to unnecessary toxicities.

OMI was used to track drug response at the single-cell level over a time course of treatment for 11 pancreatic cancer, PanIN, and ampullary cancer patients (Fig. 2). Time courses are vital to evaluate response, because cells often evolve mechanisms of drug resistance that are not immediately apparent. For example, Patient 17 organoids initially responded to 5-FU (p<0.05 vs. control, Glass’s Δ = 1.66 at 24 hours), but became resistant at day 7 (p>0.05 vs. control, Glass’s Δ = −0.04). This is a major advantage of OMI over methods that destroy the sample. Drug response was also evaluated using OMI in patient-derived fibroblasts, which grew along with organoids for most patients. The dense fibrotic extracellular matrix surrounding pancreatic tumors can hinder drug delivery by reducing blood flow and raising interstitial fluid pressure (60, 61). Thus, it may be vital to evaluate whether drugs can target both the tumor and its stromal microenvironment to enhance delivery (16, 17). For example, Patient 18’s organoids showed response to the combination of TAK-228 and ABT-263 (Glass’s Δ > 0.75) at 72 hours while co-cultured Patient 18 fibroblasts did not, suggesting that this drug regimen could successfully kill tumor cells but may have difficulty reaching them. This highlights the need for OMI-organoid technology to aid in the discovery of new treatment strategies to target both a tumor and its stroma. Additionally, there were multiple instances of novel drug combinations outperforming standard chemotherapies. For instance, TAK-228 and ABT-263 elicited a more significant metabolic response in both Patient 3 organoids and fibroblasts than the prescribed standard gemcitabine and 5-FU regimen.

Our group has shown that OMI non-invasively distinguishes unique groups of cells by their metabolic properties in human breast cancer (36, 38), human head and neck cancer (37), and murine PDAC (7). Here, we performed subpopulation analysis on Patient 13 organoid cells to evaluate whether OMI could distinguish cells with distinct drug responses in human pancreatic cancer (Fig. 3; Supplementary Fig. S6). Indeed, many treatments caused two distinct subpopulations to form in organoids within 72 hours post-treatment, including the standard chemotherapy regimen given to the patient prior to surgery and organoid generation (gemcitabine with nab-paclitaxel). This suggests that a drug-resistant cell subpopulation persisted throughout the patient’s neoadjuvant treatment, and was captured in the organoids. Accordingly, pathology of the patient’s resected tumor indicated a poor response to gemcitabine with nab-paclitaxel.

Conversely, targeted therapies induced single populations of homogenous response in Patient 13 organoids. While the combination of gemcitabine and nab-paclitaxel caused a similar average reduction in OMI index as the combination of TAK-228 and ABT-263, the former showed heterogeneity and a drug-resistant population, while the latter showed a homogeneous response. This suggests that this targeted therapy combination (TAK-228 and ABT-263) may have been a beneficial alternative for Patient 13. One month after surgery, metastasis in the liver was found on an ultrasound, and the patient died less than two weeks later. This highlights the need for a technology that can rapidly evaluate drug sensitivity in patient cells.

Untreated Patient 13 organoids were also deep-sequenced to determine if genetic heterogeneity could be the source of metabolic subpopulations in untreated organoids. Our metabolic imaging data shows greater heterogeneity than predicted by deep sequencing, as no subclonal populations with pathogenic mutations were found in these organoids. This suggests that the OMI index heterogeneity is likely not genetic in nature, and that OMI is more effective for detecting divergent populations in organoid cultures than destructive sequencing techniques. This work also highlights the importance of metabolic heterogeneity within cancer samples to capture therapeutic response. At 72 hours, a subset of Patient 13 organoids were fixed and stained for proliferation and apoptosis markers. In some cases, OMI detected shifts in metabolism with treatment that had not triggered apoptosis, suggesting that metabolic changes can precede Ki67/CC3 indicators of cell fate, and are an earlier indicator of drug efficacy.

The combination of gemcitabine and nab-paclitaxel was evaluated *in vivo* in a PDX line established from Patient 13 organoids. A small but transient response in average tumor growth was detected (Supplementary Fig. S9). This poor response is in agreement with the heterogeneous effect found in organoids using OMI. While the PDX model accurately captured the presence of drug resistance in Patient 13’s cells, it required months to first establish the PDX line, expand it, and then assess a time course of treatment. While PDX models are an important tool, our studies show that drug screens on organoids can provide more detailed response information with increased cost effectiveness in a clinically meaningful time frame.

OMI of organoids generated from tissues collected at surgery agreed with patient outcome on adjuvant therapy (Fig. 4-6). Treatment response in organoids was evaluated on the basis of both metabolic heterogeneity in treated organoids, as well as the effect size of the OMI index response. For Patients 1, 2, 3, 6, 8, and 14, OMI of organoid heterogeneity predicted whether or not the patient survived one year post-surgery without a recurrence. Based on data from this initial patient cohort, a proposed OMI index effect size cutoff of 0.75, in parallel with a decrease in wH-index could classify patients as predicted responders vs. non-responders. We cannot yet evaluate our prediction that Patient 18 will experience a recurrence within one year of surgery. Overall, our results suggest that early metabolic responses in pancreatic organoids can successfully capture the response of tumor cells *in vivo* that are not removed during surgery. In all cases, organoid viability yielded sufficient material to measure heterogeneity in response to the patient’s prescribed drug treatment and evaluate multiple alternative drug options, highlighting the potential of OMI of organoids to aid in drug discovery and development within diverse patient populations. While many other factors beyond tumor cell treatment response affect pancreatic cancer survival in the adjuvant setting (i.e. surgical margins, stage, grade, etc.), the organoid-OMI platform can be used to identify drug resistance and eliminate objectively poor drug options.

Finally, we used a dimensionality reduction approach to determine the influence of patient of origin on cellular metabolism in pancreatic organoids (Fig. 7). UMAP analysis was performed on metabolic parameters from all 45,000+ cells, completely naïve to patient of origin. These organoid cells clearly clustered by patient, but also show clear heterogeneity within each patient. This reinforces the fact that cells from a patient are metabolically heterogeneous, but more metabolically similar to each other than cells from other patients. This metabolic diversity between patients as well as within patients underlines the need for personalized medicine approaches to cancer treatment that incorporate single-cell metabolic measurements.

Organoids offer a compelling platform for the interrogation of a variety of drugs *ex vivo*. This platform provides an extracellular matrix that can support co-culture of both patient-derived cancer cells and fibroblasts, thus creating a more robust model of human disease. OMI has many benefits over existing methods to assess drug response in organoids, in part because it is completely non-destructive and it measures unique features of cell metabolism. OMI can measure response on the single-cell level to assess heterogeneity as quickly as 24 hours post-treatment and can track dynamic responses over time. We used these optical imaging tools to show that organoid drug screens can assess heterogeneous drug responses in pancreatic cancer patients (and one rare ampullary cancer patient) within a clinically meaningful time frame. Additionally, the ability to detect a drug-resistant subpopulation of cells following neoadjuvant treatment failure was demonstrated with OMI, and alternative treatment options were evaluated. In cases where multiple treatment options are likely beneficial, clinicians could use this platform to select the regimen that presents the fewest side effects for the patient, significantly improving quality of life. Taken together, OMI of organoids is a sensitive, high-throughput tool to select the best treatments for individual pancreatic cancer patients based on single-cell metabolic response, which could improve patient outcomes and enable new drug discovery.

## Acknowledgments

We thank the patients that graciously donated tissue to this study. Thank you to the Translation Science BioCore BioBank and its staff at the University of Wisconsin Carbone Cancer Center for the collection of donated tissue. Thank you to Amani Gillette and Mohammad Karim for their time, expertise, and assistance acquiring OMI images.

