## Supplemental Info for "Optical metabolic imaging of heterogeneous drug response in pancreatic cancer patient organoids"

**Supplementary Information Brief Descriptions**

Supplementary Table S1: Characteristics of patient tissue samples acquired from surgical resection of pancreatic lesions.

Supplementary Figure S1. Metabolic perturbation with cyanide validates OMI and organoid viability.

Supplementary Figure S2. Effect sizes of drug treatment on individual OMI endpoints in organoids.

Supplementary Figure S3. Effect sizes of drug treatment on individual OMI endpoints in patient-derived fibroblasts co-cultured with organoids.

Supplementary Figure S4. Significance of drug treatment effects on individual OMI endpoints in organoids.

Supplementary Figure S5. Significance of drug treatment effects on individual OMI endpoints in patient-derived fibroblasts co-cultured with organoids.

Supplementary Figure S6. Dual immunofluorescence of cell fate in pancreatic organoids.

Supplementary Figure S7. Correlation between OMI index, proliferation, and apoptosis in pancreatic organoids.

Supplementary Figure S8. Growth response to gemcitabine and nab-paclitaxel combination therapy in tumors grown from Patient 13 organoids in athymic nude mice.

Supplementary Figure S9. Response to patient adjuvant therapy in patient-derived fibroblasts.

**Supplementary Table S1**
**Characteristics of patient tissue samples acquired from surgical resection of pancreatic lesions. RFS, recurrence-free survival. Asterisk indicates that a recurrence has occurred.**

| Patient # | Drug treatment before surgery | Neoadjuvant Treatment Response | Post-treatment Diagnosis | Residual Tumor Size | Stage | Viable Organ-oids | Drug treatment after surgery | Known RFS (months) |
|---|---|---|---|---|---|---|---|---|
| 1 | 5-FU | Partial response, score 2 | PDAC, well-differentiated | 3.5x1.8x1.7 cm | ypT3N0 | Yes | Gemcitabine + 5-FU | >24 |
| 2 | None | N/A | PDAC, poorly-differentiated | 4.8x3.5x2 cm | pT3N1 | Yes | Gemcitabine + 5-FU | 14.5* |
| 3 | 5-FU | Poor or no response, score 3 | PDAC, poorly-differentiated | 2x2x1.5 cm | ypT3N1 | Yes | Gemcitabine + 5-FU | 11.5* |
| 4 | None | N/A | Mucinous adenocarcinoma | 4x3x2.9 cm | pT3N1 | No | Gemcitabine + 5-FU | >17 |
| 5 | FOLFIRINOX | Complete response, score 0 | No residual cancer | N/A | ypT0N0 | Yes | None | N/A |
| 6 | None | N/A | Ampullary adenocarcinoma, moderately-differentiated | 2x2x1.2 cm | pT2N1 | Yes | Oxaliplatin + 5-FU | >13 |
| 7 | None | N/A | PDAC, moderately-differentiated (arising in IPMN) | 0.5 cm | pT1N0 | No | None | >14 |
| 8 | Gemcitabine + nab-paclitaxel | Partial response, score 2 | PDAC, moderately-differentiated | 3.0 cm | ypT3N1 | Yes | 5-FU | 5* |
| 9 | Gemcitabine + nab-paclitaxel + FOLFIRINOX | Poor or no response, score 3 | PDAC, moderately-differentiated | 3.1x2.5x2.3 cm | ypT4N1 | No | None | 4* |
| 10 | None | N/A | PDAC, moderately-differentiated | 3.5x2x2 cm | pT3N1 | No | Unknown | 2* |
| 11 | None | N/A | Chronic pancreatitis and PanIN-1 | N/A | N/A | Yes | None | N/A |
| 12 | None | N/A | PDAC, poorly-differentiated | 1.2x1.1x1.1 cm | pT1N1 | Yes | FOLFIRINOX | 2* |
| 13 | Gemcitabine + nab-paclitaxel | Poor or no response, score 3 | Undifferentiated (anaplastic) carcinoma of pancreas | 4.9x4.3x4.1 cm | ypT3N1 | Yes | None | 0.5* |
| 14 | None | N/A | PDAC, moderately-differentiated | 3.5 cm | pT3pN1 | Yes | Gemcitabine + 5-FU | >13 |
| 15 | None | N/A | PDAC, moderately-differentiated | 3x2.5x2.5 cm | pT2N1 | No | Gemcitabine + 5-FU | >7 |
| 16 | Gemcitabine + nab-paclitaxel | Partial response, score 2 | Ampullary adenocarcinoma, well-differentiated | 2.5x2.5x1.4 cm | ypT3bN1 | No | None | >3 |
| 17 | FOLFIRINOX | Partial response, score 2 | PDAC, well-differentiated | 2.0x2.0x1.5 cm | ypT2N0 | Yes | None | 4* |
| 18 | None | N/A | PDAC, poorly-differentiated | 4.7x4.3x3.5 cm | pT3N2 | Yes | FOLFIRINOX | >2 |

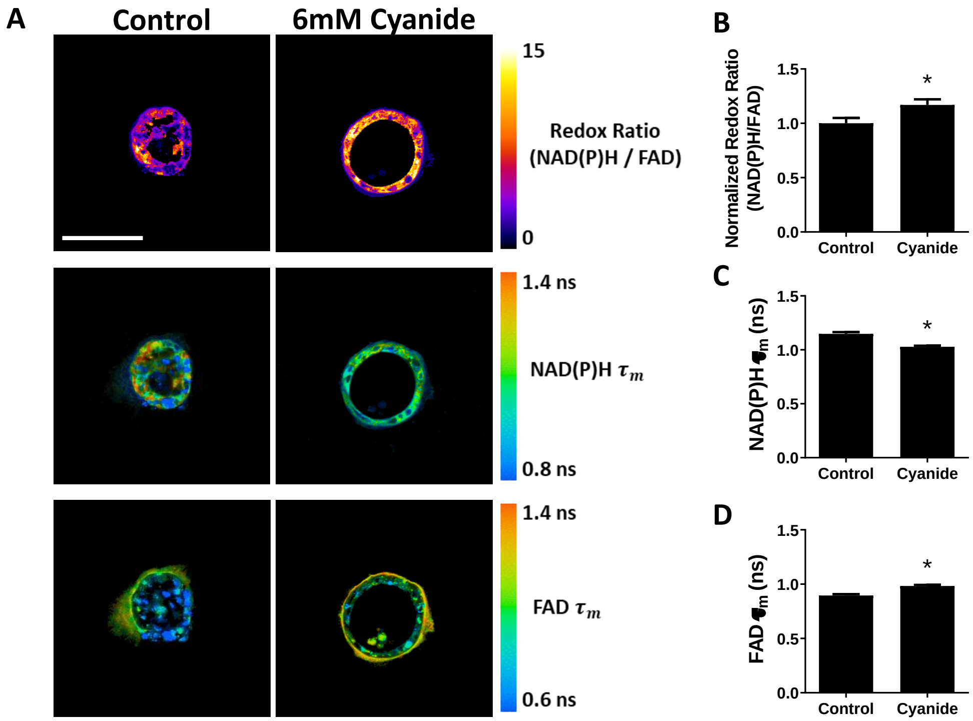


**Supplementary Figure S1. Metabolic perturbation with cyanide validates OMI and organoid viability. A,** Representative redox ratio, NAD(P)H $\tau_m$, and FAD $\tau_m$ images of Patient 1 pancreatic organoids before and after 6mM cyanide treatment. Scale bar is 100 µm. **B,** Optical redox ratio increases with cyanide. **C,** NAD(P)H mean lifetime decreases with cyanide. **D,** FAD mean lifetime increases with cyanide treatment. Error bars indicate mean ± SD. * $p<0.05$. N=4 pancreatic organoids comprising 64 cells.

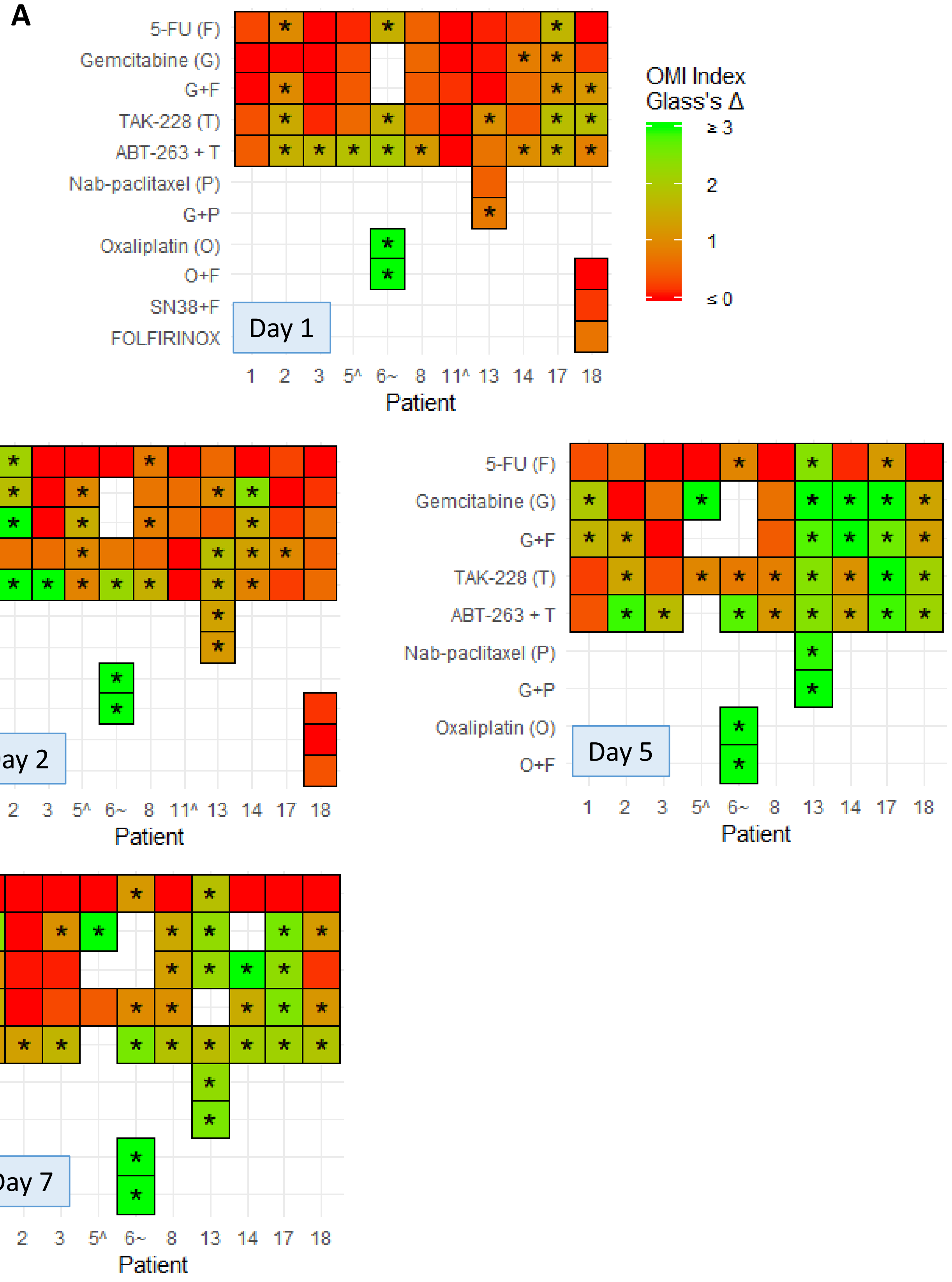

A
5-FU (F)
Gemcitabine (G)
G+F
TAK-228 (T)
ABT-263 + T
Nab-paclitaxel (P)
G+P
Oxaliplatin (O)
O+F
SN38+F
FOLFIRINOX
Day 1
1 2 3 5^ 6~ 8 11^ 13 14 17 18
Patient
OMI Index
Glass's Δ
≥ 3
2
1
≤ 0
Day 2
Day 5
Day 7

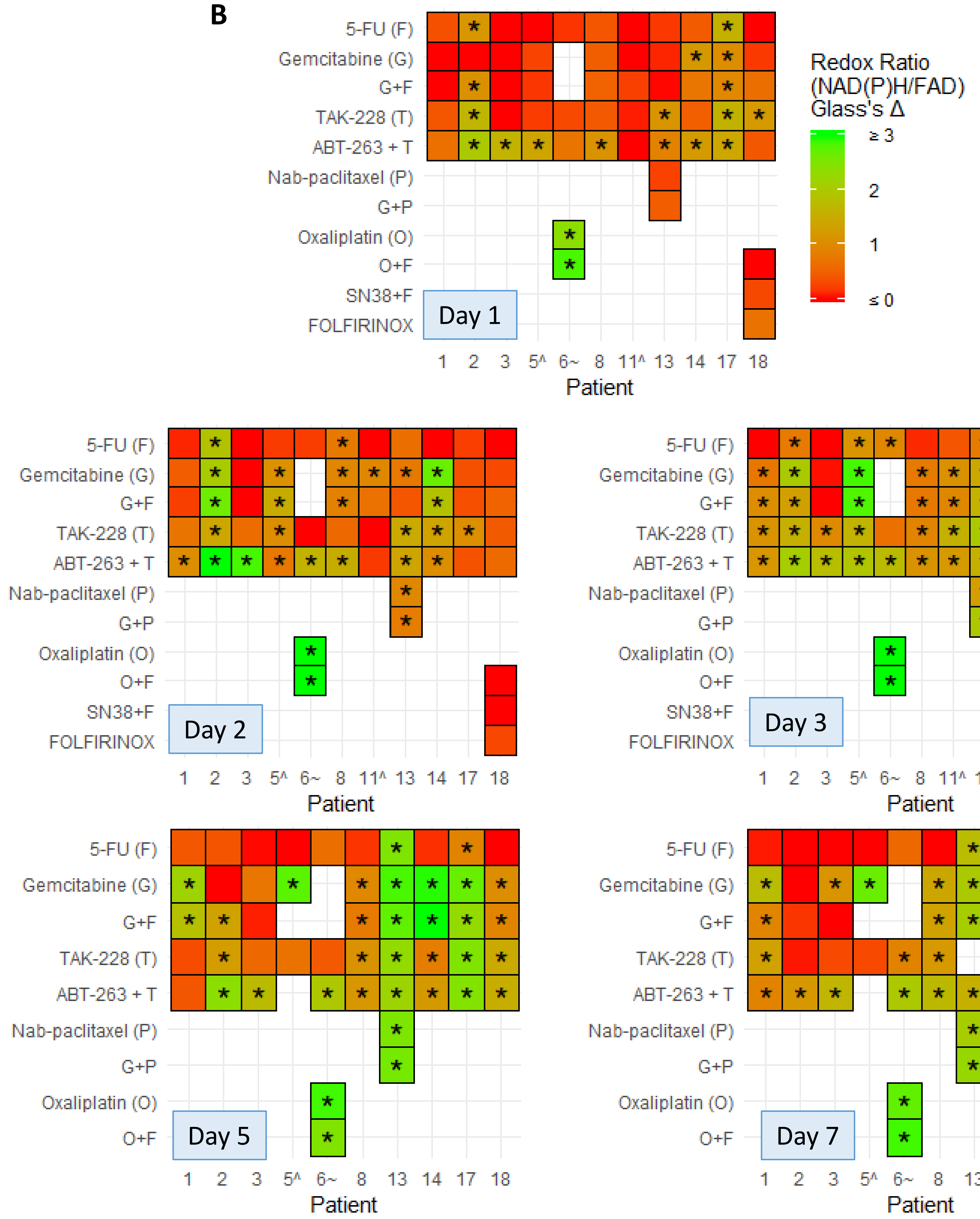
B
Redox Ratio (NAD(P)H/FAD) Glass's Δ
≥ 3
2
1
≤ 0
5-FU (F)
Gemcitabine (G)
G+F
TAK-228 (T)
ABT-263 + T
Nab-paclitaxel (P)
G+P
Oxaliplatin (O)
O+F
SN38+F
FOLFIRINOX
Day 1
Day 2
Day 3
Day 5
Day 7
1 2 3 5^ 6~ 8 11^ 13 14 17 18
1 2 3 5^ 6~ 8 13 14 17 18
Patient

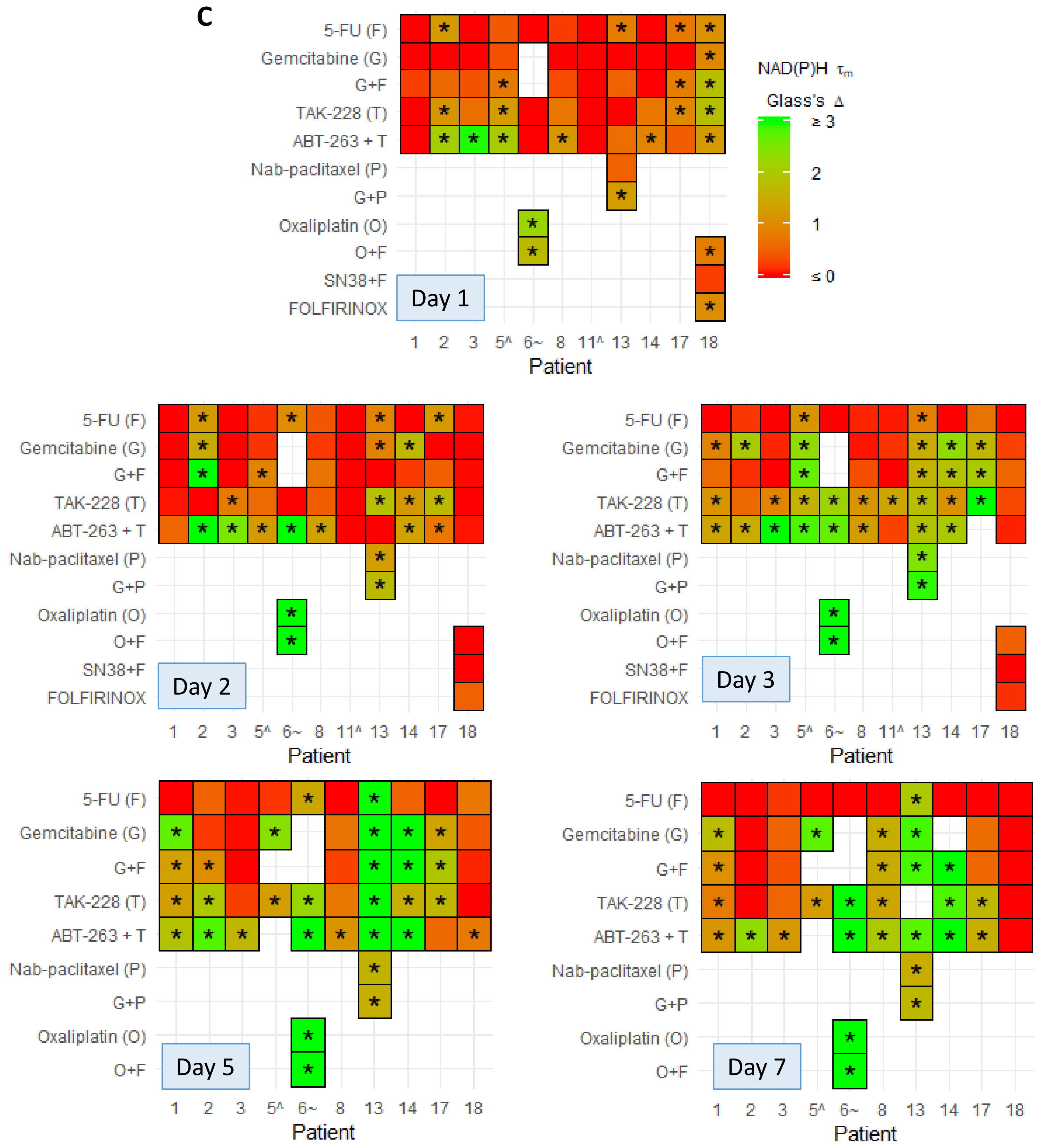
C
NAD(P)H τm
Glass's Δ
≥ 3
2
1
≤ 0
5-FU (F)
Gemcitabine (G)
G+F
TAK-228 (T)
ABT-263 + T
Nab-paclitaxel (P)
G+P
Oxaliplatin (O)
O+F
SN38+F
FOLFIRINOX
Day 1
Day 2
Day 3
Day 5
Day 7
1 2 3 5^ 6~ 8 11^ 13 14 17 18
Patient

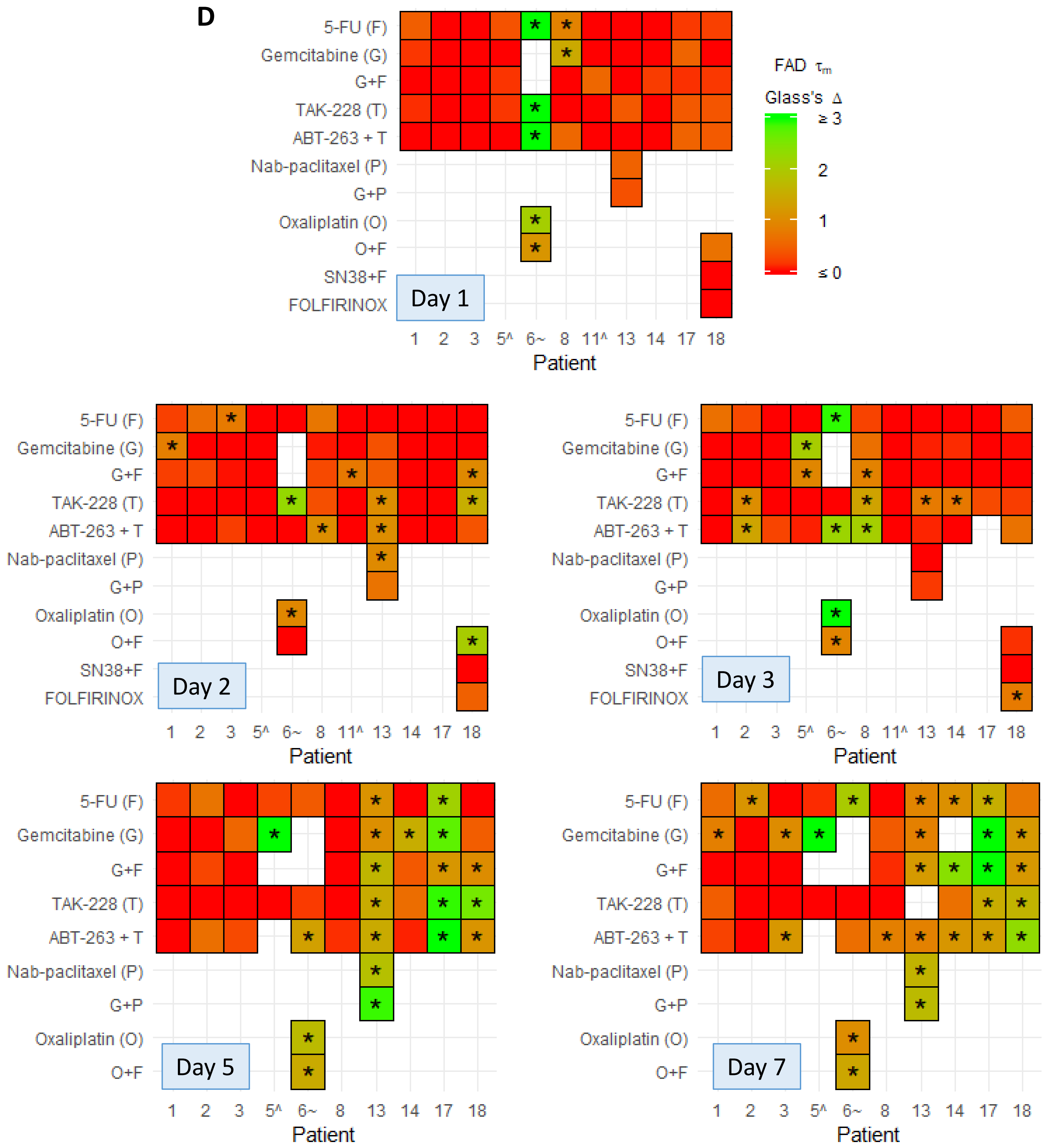

D
5-FU (F)
Gemcitabine (G)
G+F
TAK-228 (T)
ABT-263 + T
Nab-paclitaxel (P)
G+P
Oxaliplatin (O)
O+F
SN38+F
FOLFIRINOX
Day 1
Day 2
Day 3
Day 5
Day 7
1 2 3 5^ 6~ 8 11^ 13 14 17 18
Patient
FAD τm
Glass's Δ
≥ 3
2
1
≤ 0

**Supplementary Figure S2. Effect sizes of drug treatment on individual OMI endpoints in organoids. A-D,** Heatmap representation of the treatment effect size (Glass's Δ) at each time point for the OMI index (**A**), redox ratio (**B**), NAD(P)H $\tau_m$ **(C)**, and FAD $\tau_m$ (**D**). '^' indicates the patient lesion was diagnosed as PanIN. '~' indicates the patient lesion was diagnosed as ampullary cancer. * Glass's Δ ≥ 0.75 vs. control.

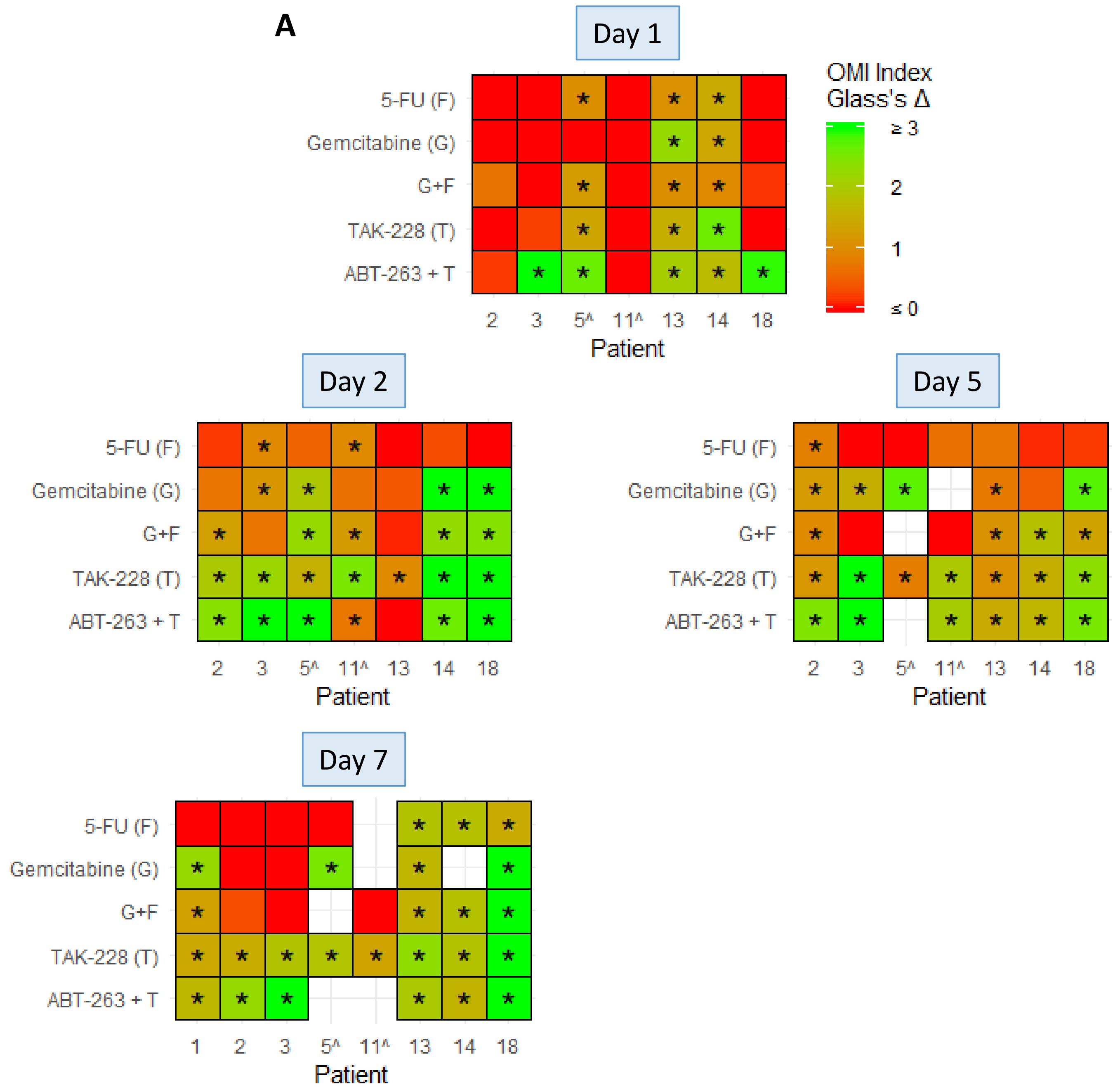
A
Day 1
OMI Index
Glass's Δ
≥ 3
2
1
≤ 0
5-FU (F)
Gemcitabine (G)
G+F
TAK-228 (T)
ABT-263 + T
2 3 5^ 11^ 13 14 18
Patient
Day 2
Day 5
Day 7
1 2 3 5^ 11^ 13 14 18
Patient

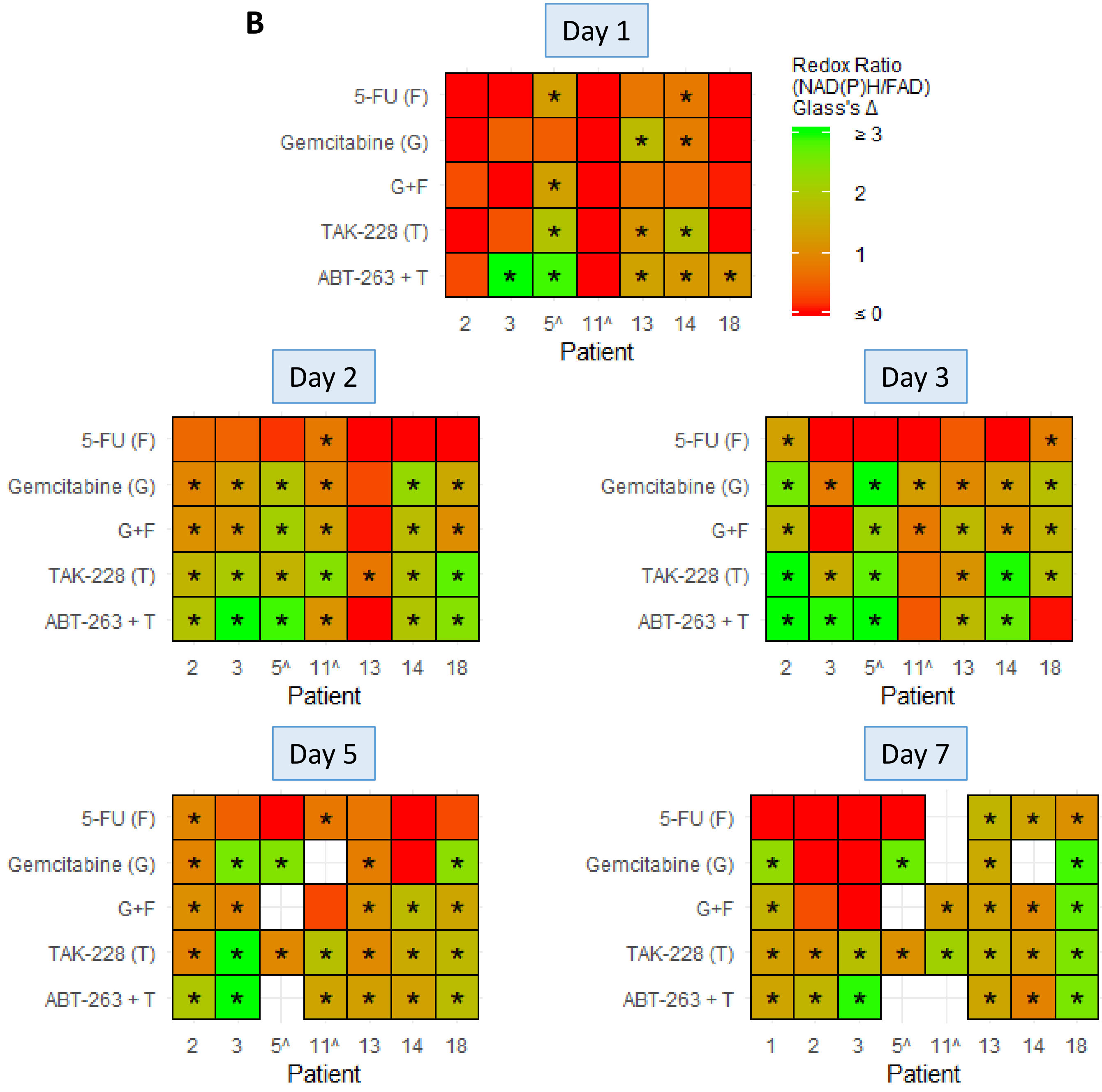
B
Day 1
Day 2
Day 3
Day 5
Day 7
5-FU (F)
Gemcitabine (G)
G+F
TAK-228 (T)
ABT-263 + T
1
2
3
5^
11^
13
14
18
Patient
Redox Ratio
(NAD(P)H/FAD)
Glass's Δ
≥ 3
2
1
≤ 0

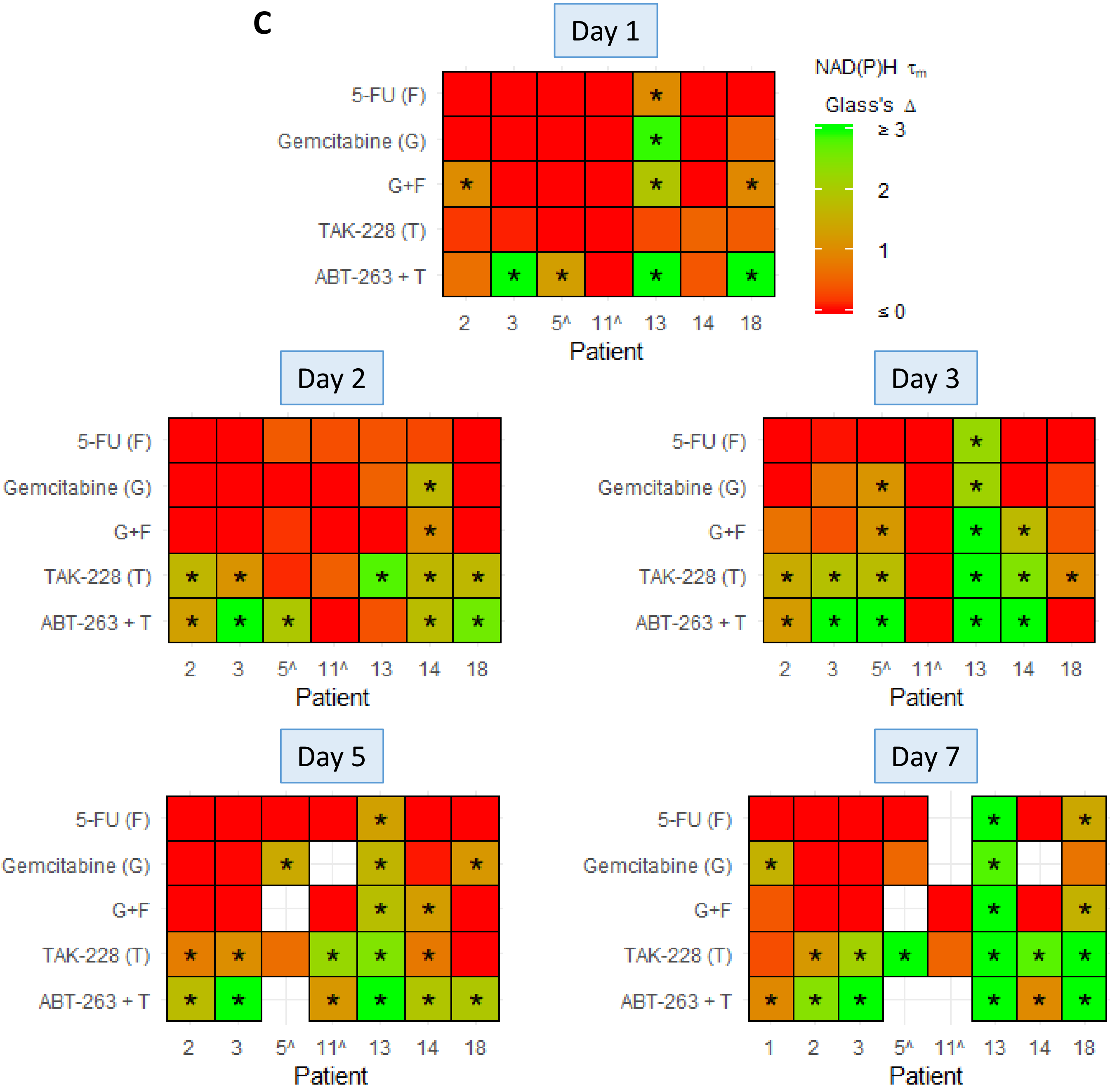
C
Day 1
Day 2
Day 3
Day 5
Day 7
NAD(P)H $\tau_m$
Glass's Δ
≥ 3
2
1
≤ 0
5-FU (F)
Gemcitabine (G)
G+F
TAK-228 (T)
ABT-263 + T
Patient
1
2
3
5^
11^
13
14
18

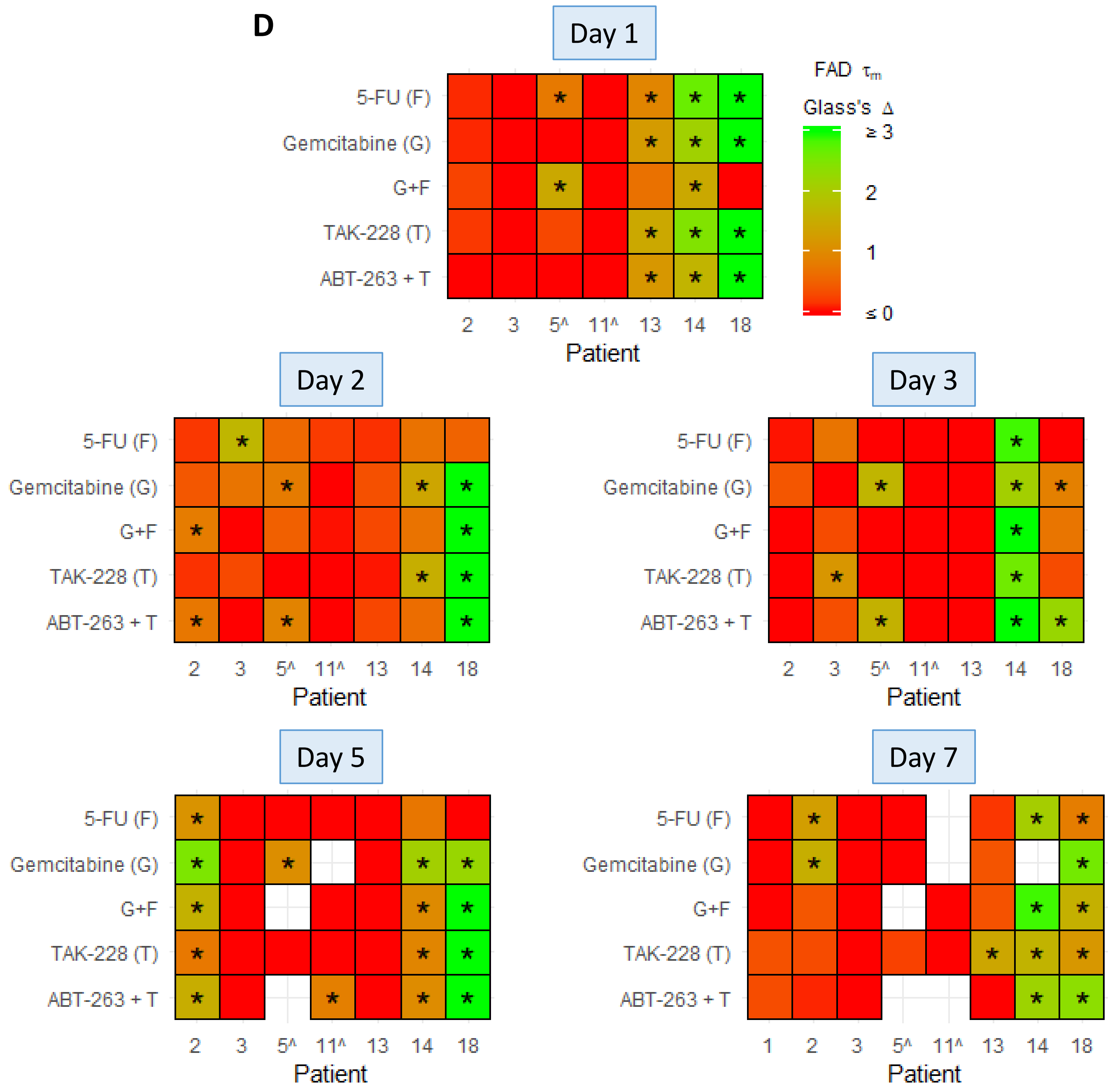


**Supplementary Figure S3. Effect sizes of drug treatment on individual OMI endpoints in patient-derived fibroblasts co-cultured with organoids. A-D,** Heatmap representation of the treatment effect size (Glass's Δ) at each time point for the OMI index (**A**), redox ratio (**B**), NAD(P)H $\tau_m$ **(C)**, and FAD $\tau_m$ (**D**). '^' indicates the patient lesion was diagnosed as PanIN. '~' indicates the patient lesion was diagnosed as ampullary cancer. * Glass's $\Delta \geq 0.75$ vs. control.

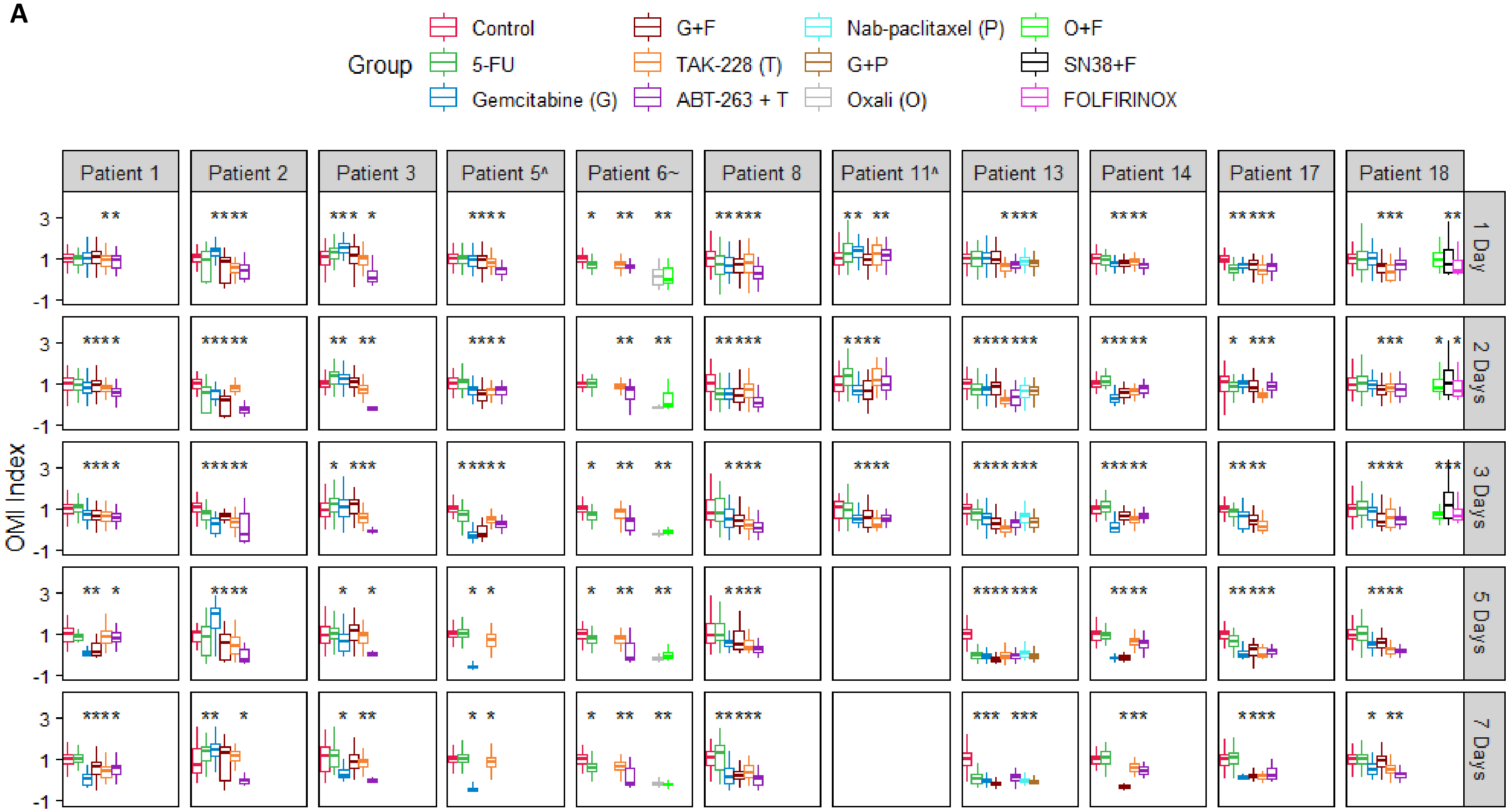
A
Group
Control
5-FU
Gemcitabine (G)
G+F
TAK-228 (T)
ABT-263 + T
Nab-paclitaxel (P)
G+P
Oxali (O)
O+F
SN38+F
FOLFIRINOX
Patient 1
Patient 2
Patient 3
Patient 5^
Patient 6~
Patient 8
Patient 11^
Patient 13
Patient 14
Patient 17
Patient 18
1 Day
2 Days
3 Days
5 Days
7 Days
OMI Index

**B**

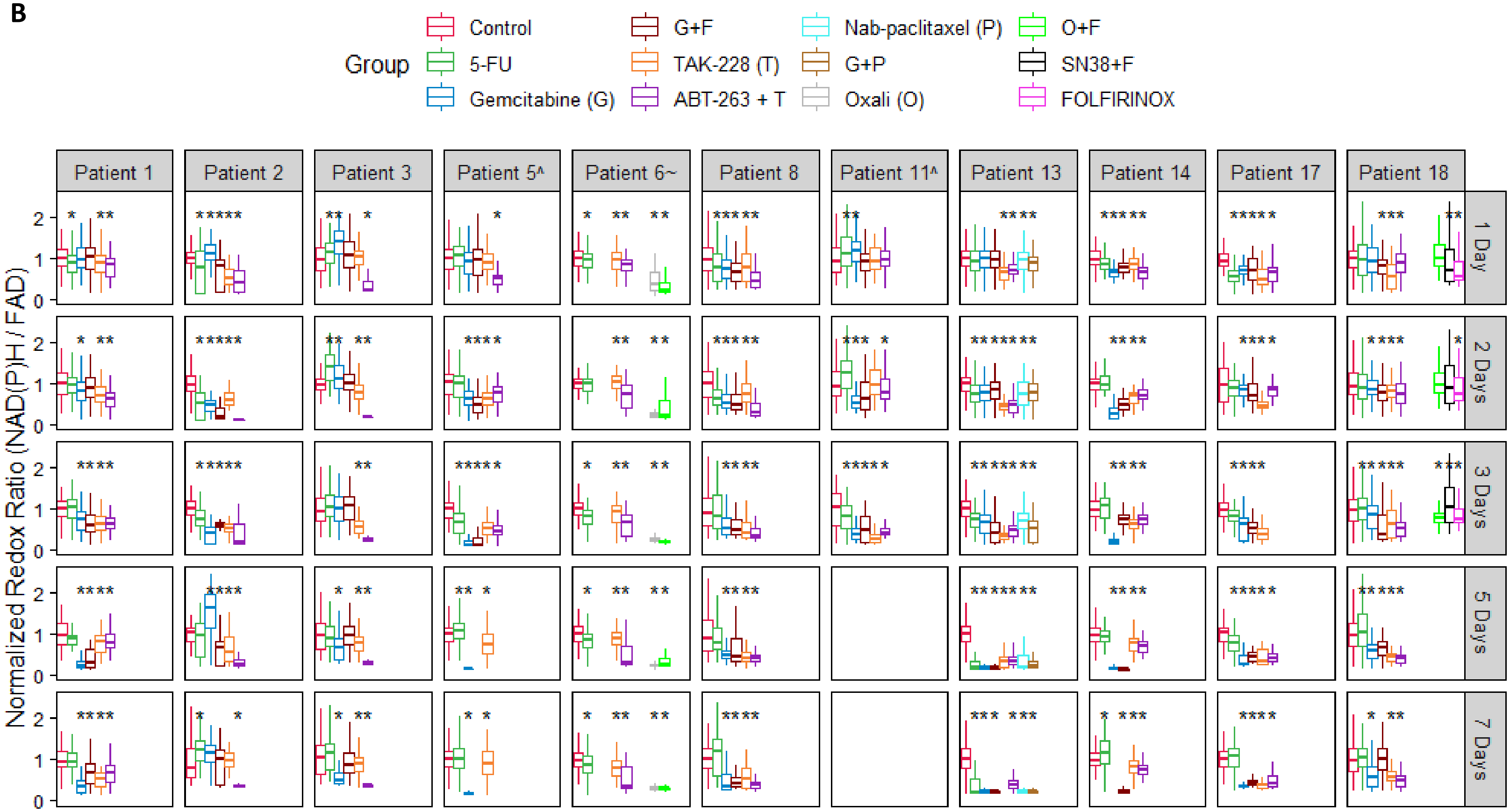

Group
Control
5-FU
Gemcitabine (G)
G+F
TAK-228 (T)
ABT-263 + T
Nab-paclitaxel (P)
G+P
Oxali (O)
O+F
SN38+F
FOLFIRINOX
Patient 1
Patient 2
Patient 3
Patient 5^
Patient 6~
Patient 8
Patient 11^
Patient 13
Patient 14
Patient 17
Patient 18
1 Day
2 Days
3 Days
5 Days
7 Days
Normalized Redox Ratio (NAD(P)H / FAD)
0
1
2

C

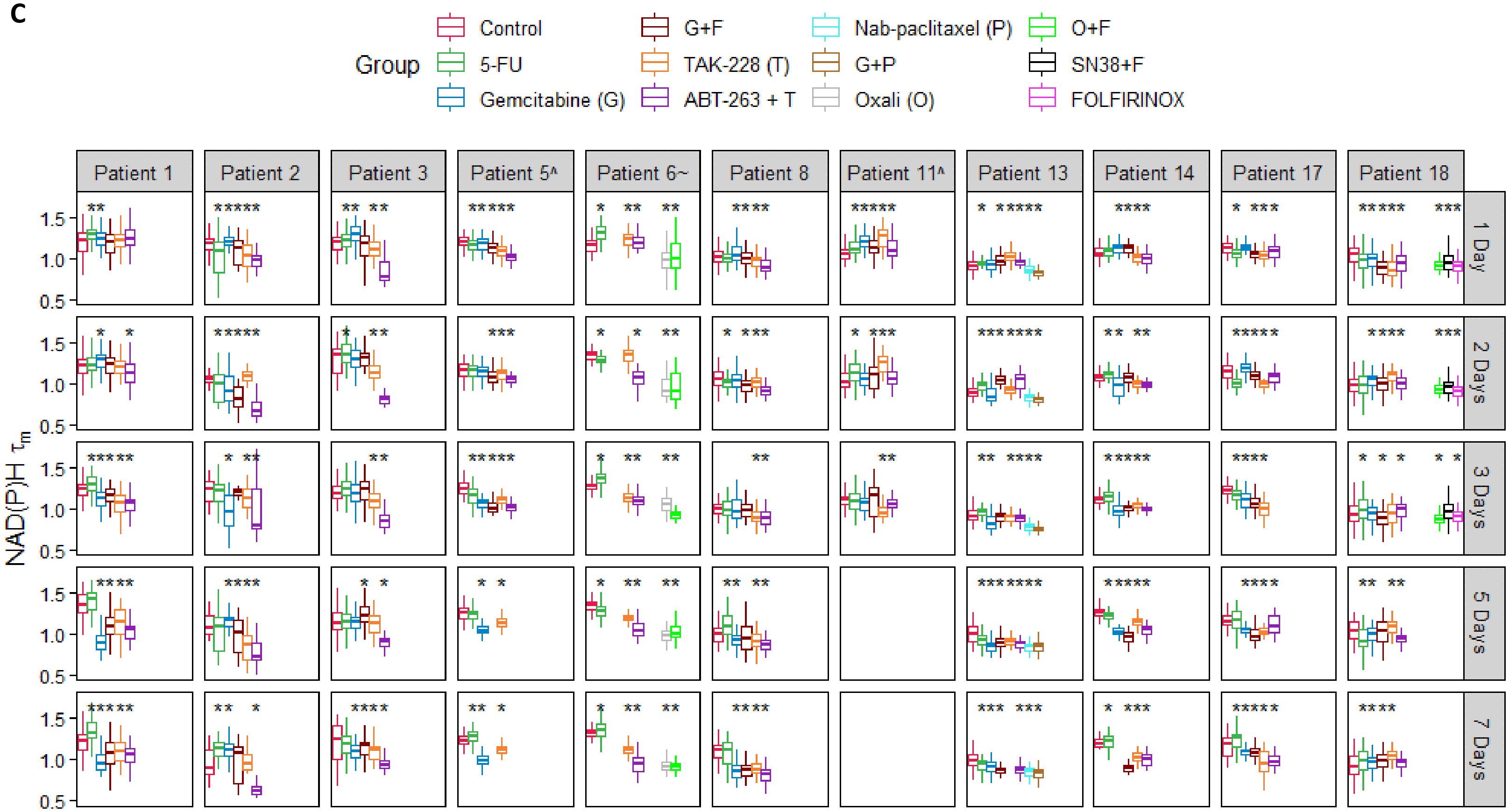

Group
Control
5-FU
Gemcitabine (G)
G+F
TAK-228 (T)
ABT-263 + T
Nab-paclitaxel (P)
G+P
Oxali (O)
O+F
SN38+F
FOLFIRINOX
Patient 1
Patient 2
Patient 3
Patient 5^
Patient 6~
Patient 8
Patient 11^
Patient 13
Patient 14
Patient 17
Patient 18
1 Day
2 Days
3 Days
5 Days
7 Days
NAD(P)H $\tau_m$
0.5
1.0
1.5

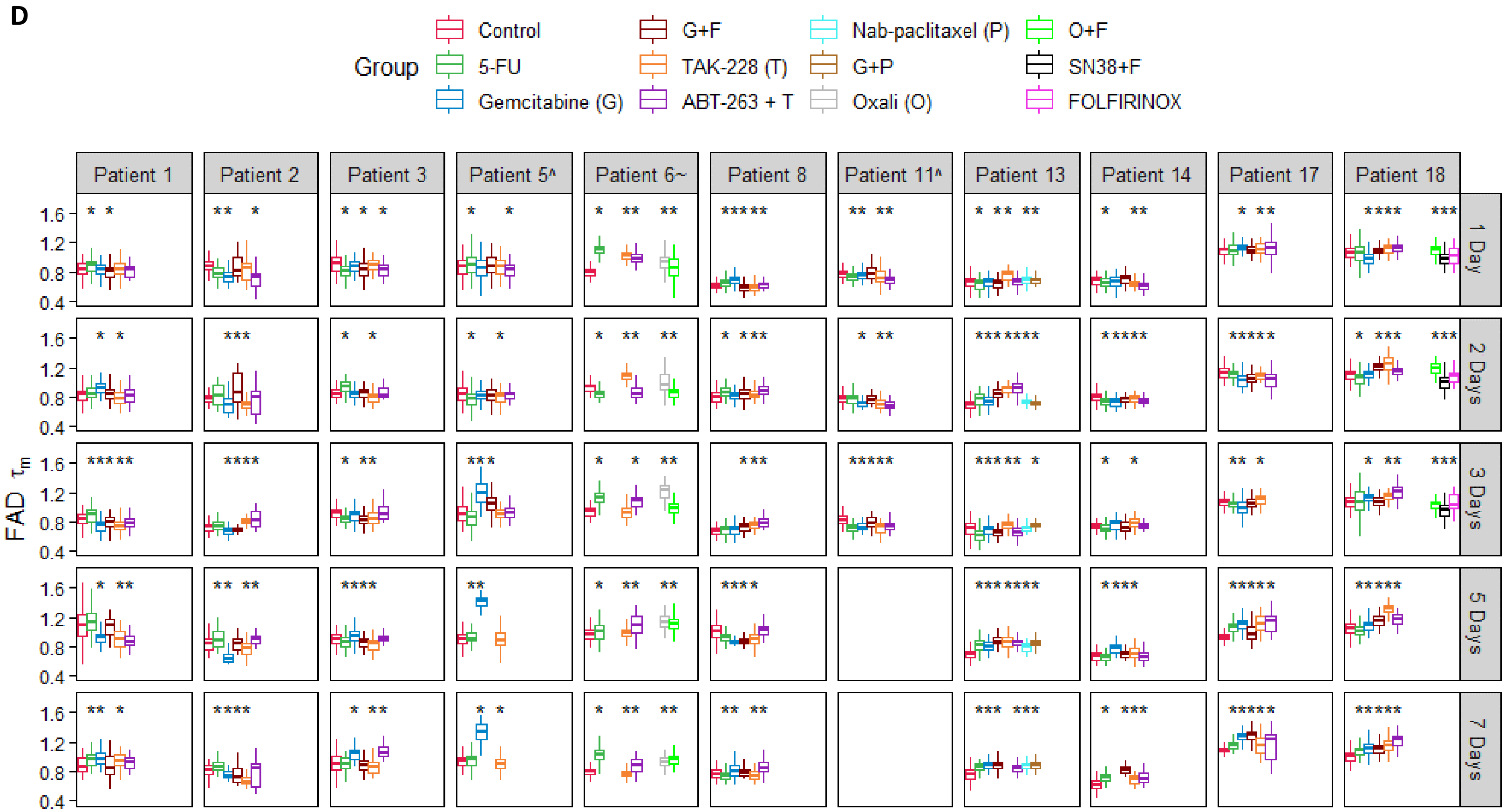


**Supplementary Figure S4. Significance of drug treatment effects on individual OMI endpoints in organoids. A-D,** Boxplot summaries comparing the effect of all drugs between patients and time points in organoids on the OMI index (**A**), redox ratio (**B**), NAD(P)H $\tau_m$ (**C**), and FAD $\tau_m$ (**D**). '^' indicates the patient lesion was diagnosed as PanIN. '~' indicates the patient lesion was diagnosed as ampullary cancer. * p<0.05 vs. control.

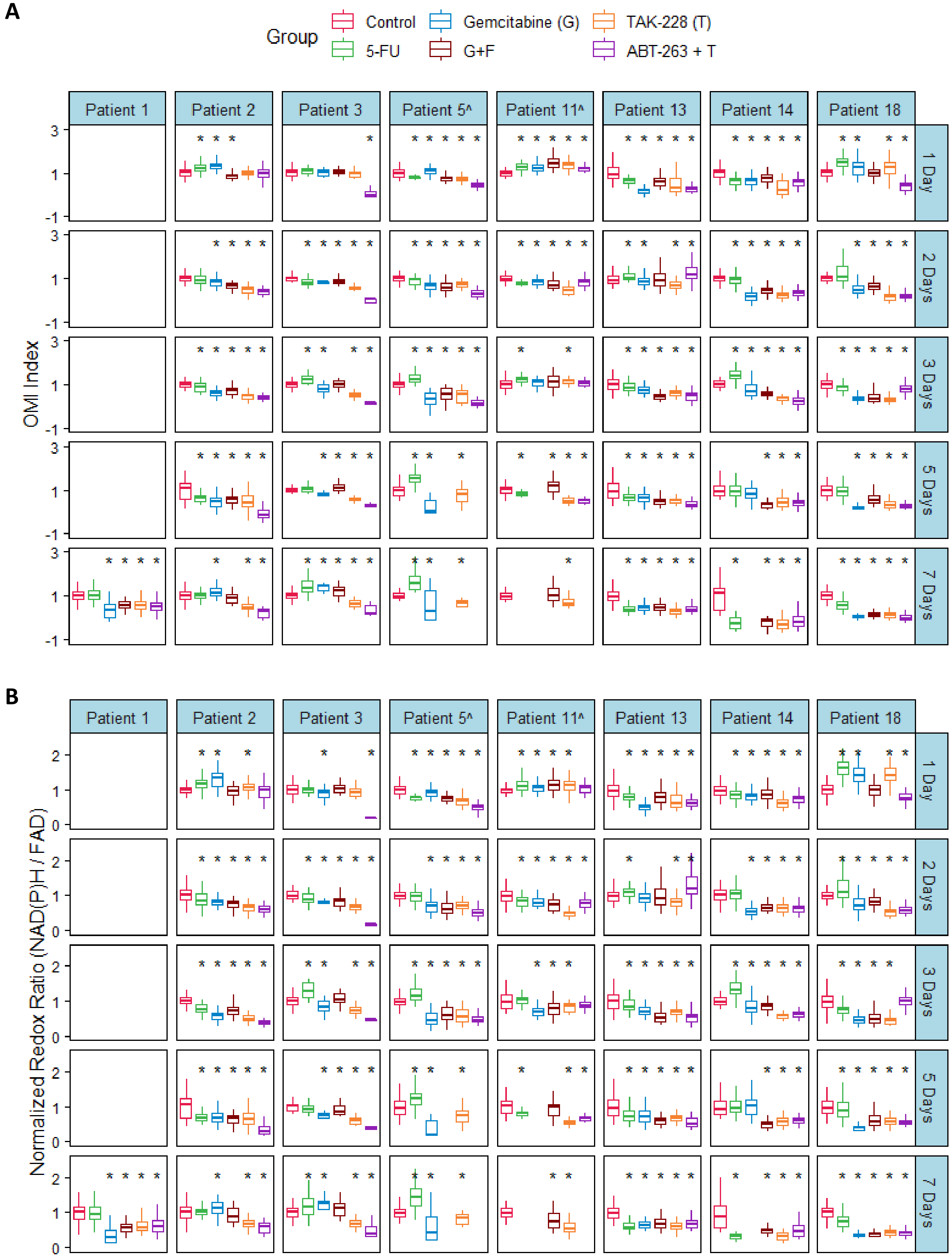
A
Group
Control
Gemcitabine (G)
TAK-228 (T)
5-FU
G+F
ABT-263 + T
Patient 1
Patient 2
Patient 3
Patient 5^
Patient 11^
Patient 13
Patient 14
Patient 18
1 Day
2 Days
3 Days
5 Days
7 Days
OMI Index
B
Normalized Redox Ratio (NAD(P)H / FAD)

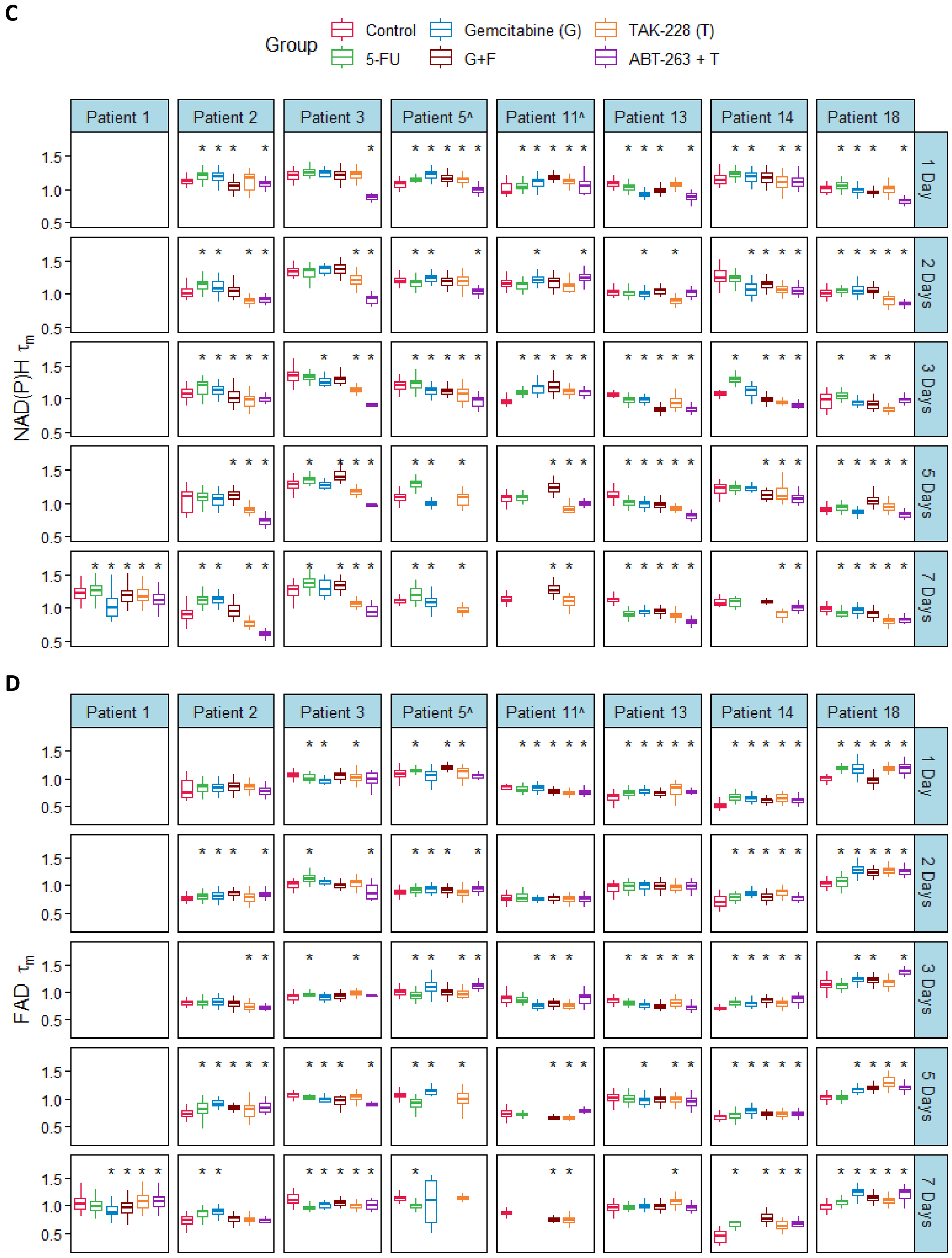

C
Group
Control
5-FU
Gemcitabine (G)
G+F
TAK-228 (T)
ABT-263 + T
Patient 1
Patient 2
Patient 3
Patient 5^
Patient 11^
Patient 13
Patient 14
Patient 18
1 Day
2 Days
3 Days
5 Days
7 Days
NAD(P)H τm
D
FAD τm

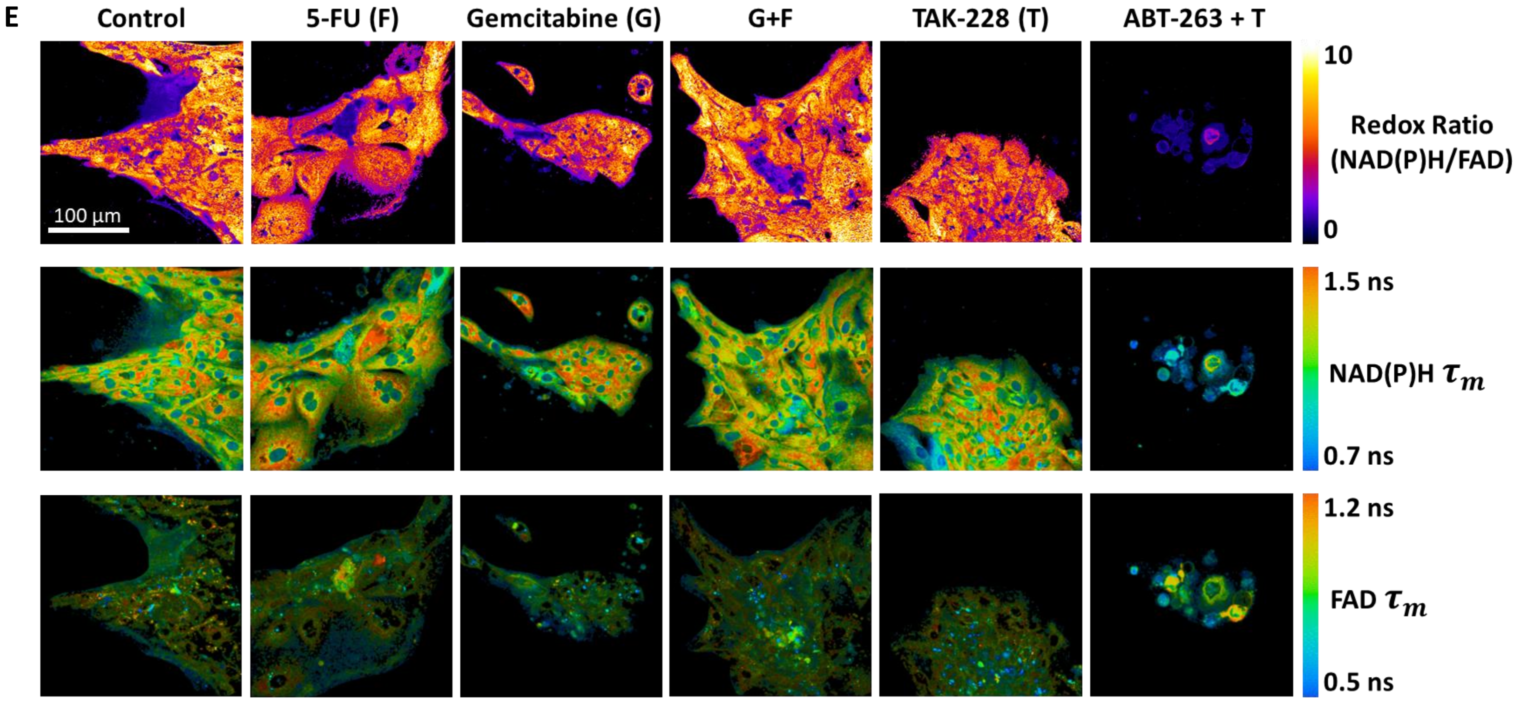


**Supplementary Figure S5. Significance of drug treatment effects on individual OMI endpoints in patient-derived fibroblasts co-cultured with organoids. A-D,** Boxplot summaries comparing the effect of all drugs on patient-derived fibroblasts at all time points for the OMI index (**A**), redox ratio (**B**), NAD(P)H $\tau_m$ (**C**), and FAD $\tau_m$ (**D**). '^' indicates the patient lesion was diagnosed as PanIN. '~' indicates the patient lesion was diagnosed as ampullary cancer. * $p<0.05$ vs. control. **E,** Representative OMI images of fibroblast monolayer co-cultured with organoids derived from Patient 3 at 24 hours of treatment.

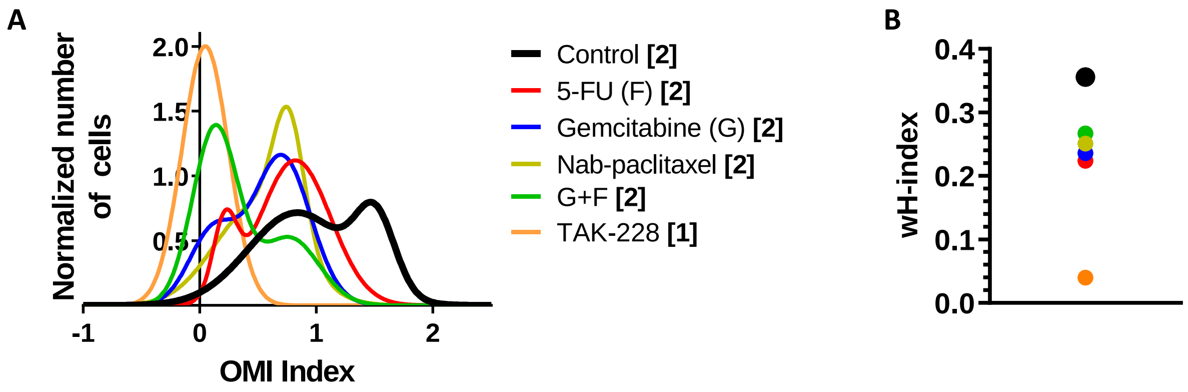


**Supplementary Figure S6. Population distribution modeling and wH-index in response to additional drugs in Patient 13 organoids. A,** Normalized density distributions of the OMI index of individual cells after 72 hours. Bracketed number indicates number of subpopulations. **B,** The effect of treatment on the wH-index of OMI index density distributions in (A). Error bars not visible. N=1000 fits/group.

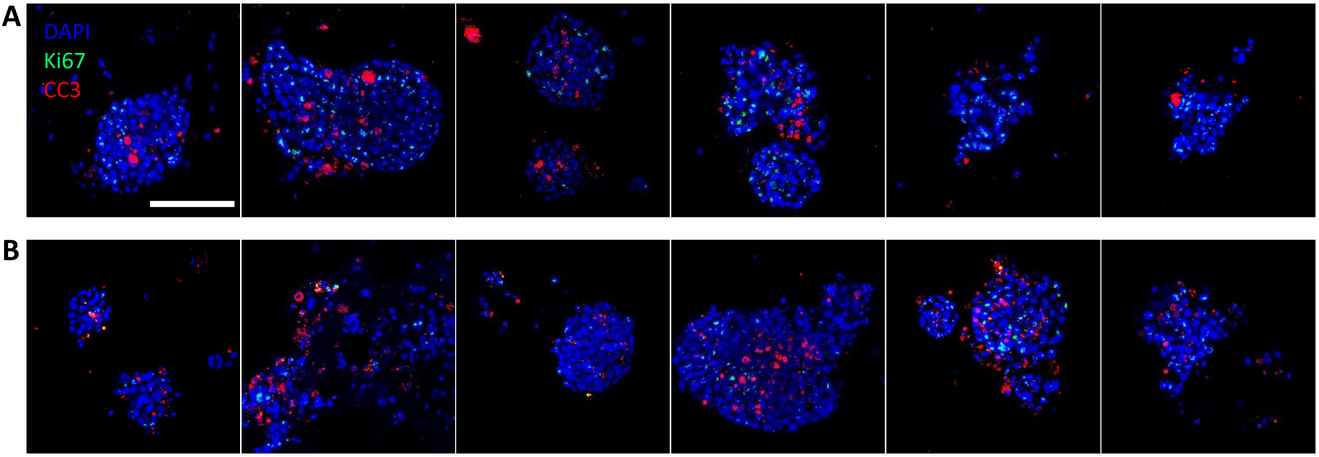


**Supplementary Figure S7. Dual immunofluorescence of cell fate in pancreatic organoids.** Composite images of Patient 13 organoids stained for Ki67 (green), cleaved caspase-3 (red), and DAPI (blue). **A,** Control organoids. Scale bar is 100 μm. **B,** organoids treated for 72 hours with the combination of TAK-228 and ABT-263.

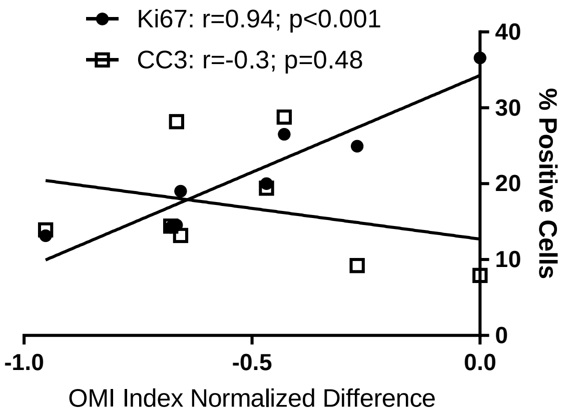


**Supplementary Figure S8. Correlation between OMI index, proliferation, and apoptosis in pancreatic organoids**. OMI index positively correlates with percentage of cells expressing Ki67 (filled circles). OMI index does not significantly correlate with percentage of cells expressing CC3 (open squares). Each dot represents the average value for one treatment group of Patient 13 organoids at 72 hours of treatment. N=8 conditions.

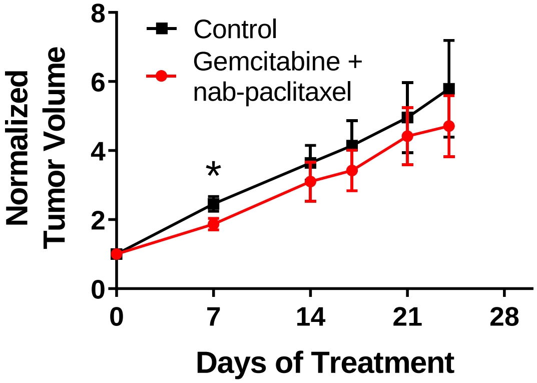


**Supplementary Figure S9. Growth response to gemcitabine and nab-paclitaxel combination therapy in tumors grown from Patient 13 organoids in athymic nude mice.** Error bars indicate mean ± SEM. * $p<0.05$ vs. control. N>20 tumors per group at day 7, N>13 tumors per group at day 14 and beyond.

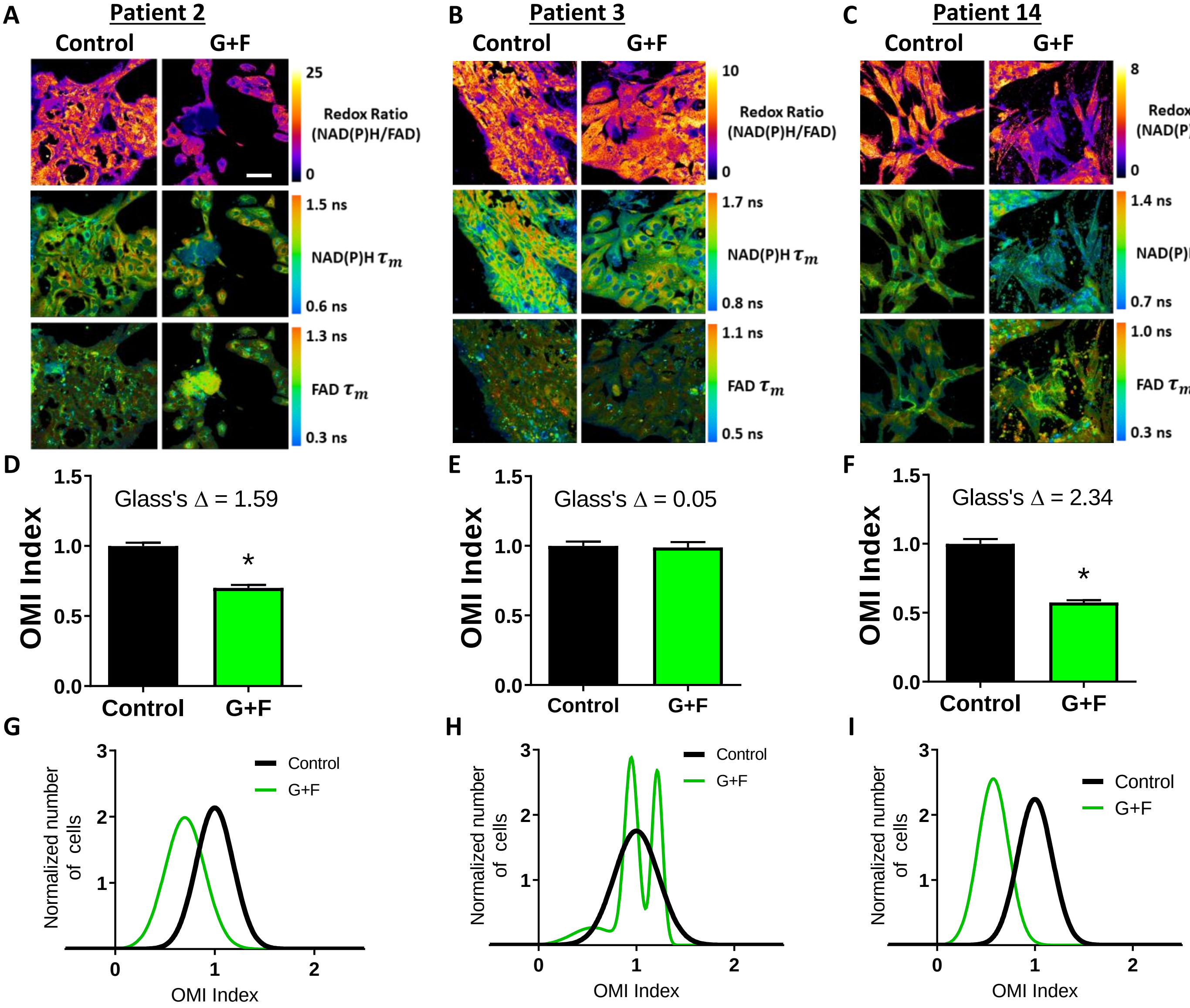


**Supplementary Figure S10. Response to patient adjuvant therapy in patient-derived fibroblasts. A-C,** Representative redox ratio, NAD(P)H $\tau_m$, and FAD $\tau_m$ images of fibroblasts co-cultured with organoids from Patients 2 (**A**), 3 (**B**), and 14 (**C**). Left columns indicate control fibroblasts, and right columns indicate fibroblasts treated with drugs matched to the patient's adjuvant treatment. G+F = gemcitabine + 5-FU. Scale bar is 50 μm. **D-F,** The effect of the same drugs on the OMI index averaged across all fibroblasts derived from Patient 2 (**D**), 3 (**E**), and 14 (**F**). Error bars indicate mean ± SEM. * p<0.0001. **G-I,** Single-cell OMI index subpopulation analysis of treatment response in fibroblasts from Patient 2 (**G**), 3 (**H**), and 14 (**I**).
